# An amphiregulin reporter mouse enables transcriptional and clonal expansion analysis of reparative lung Treg cells

**DOI:** 10.1101/2024.09.26.615245

**Authors:** Lucas F. Loffredo, Katherine A. Kaiser, Adam Kornberg, Samhita Rao, Kenia de los Santos-Alexis, Arnold Han, Nicholas Arpaia

## Abstract

Regulatory T (Treg) cells are known to play critical roles in tissue repair via provision of growth factors such as amphiregulin (Areg). Areg-producing Treg cells have previously been difficult to study because of an inability to isolate live Areg-producing cells. In this report, we created a novel reporter mouse to detect Areg expression in live cells (*Areg^Thy1.1^*). We employed influenza A and bleomycin models of lung damage to sort Areg-producing and –non-producing Treg cells for transcriptomic analyses. Single cell RNA-seq revealed distinct subpopulations of Treg cells and allowed transcriptomic comparisons of damage-induced populations. Single cell TCR sequencing showed that Treg cell clonal expansion is biased towards Areg-producing Treg cells, and largely occurs within damage-induced subgroups. Gene module analysis revealed functional divergence of Treg cells into immunosuppression-oriented and tissue repair–oriented groups, leading to identification of candidate receptors for induction of repair activity in Treg cells. We tested these using an ex vivo assay for Treg cell–mediated tissue repair, identifying 4-1BB agonism as a novel mechanism for reparative activity induction. Overall, we demonstrate that the *Areg^Thy1.1^*mouse is a promising tool for investigating tissue repair activity in leukocytes.

## Introduction

The immune response in damaged tissue must be tightly coordinated to effectively clear pathogens while allowing for proper resolution and repair. For the latter to occur correctly, immune activity must be dampened following pathogen removal to prevent further damage by the immune system itself. Regulatory T (Treg) cells are chiefly known for their roles as critical mediators of this immunosuppression [1]; however, recent studies have identified additional roles for Treg cells in tissue repair – as sources of direct signals to non-hematopoietic cells that regulate reparative processes. Several Treg cell–derived factors targeting different types of tissue cells have been identified to this effect [2], with the most studied being the EGFR ligand–family growth factor amphiregulin (Areg). The reparative effects of Areg production by Treg cells have been shown in multiple damaged tissue environments, including muscle [3], brain [4], heart [5], and lung [6].

Since the discovery of the pro-repair function of Treg cells, several questions have arisen as to the nature of Treg cells serving this role. While studies have pointed to a context-dependent induction of reparative functionality induced by alarmins such as IL-33 or inflammatory mediators such as IL-18 [6], whether reparative Treg cells represent a stable ontogenetically separate subtype from immunosuppression-oriented Treg cells is still unknown. Further, although Treg cell clones found to be expanded in adipose tissue and damaged muscle have been characterized for their tissue homing and repair capacity [3, 7], it has yet to be definitively determined if reparative Treg cells derive from uniquely clonally expanded Treg cells in response to T cell receptor (TCR) activation in damaged tissues. Importantly, the identification of novel pathways capable of inducing the pro-repair phenotype of Treg cells could be therapeutically leveraged to limit tissue damage in various sterile and pathogenic contexts.

Separate from these lines of investigation, there is a more general knowledge gap regarding T cell clonal expansion in response to non-pathogenic, sterile organ damage. While clonal expansion/effector activity towards pathogenic agents in tissue is at the core of our understanding of the adaptive immune response, this paradigm does not address the activity of T cells in damaged tissue in the absence of a pathogen. In many of these scenarios, T cells are recruited to and rapidly proliferate at the site of damage [8]. This includes Treg cells, which are greatly expanded in response to non-pathogenic damage in muscle, brain, heart, skin, and liver [3–5, 9, 10]. While expansion of Treg cells in the lung has mostly been studied in the context of pathogenic damage (e.g., influenza virus) or scenarios mimicking pathogen exposure (e.g., LPS), this increased prevalence has also been shown to occur in models of sterile lung damage, such as bleomycin exposure [11]. Whether Treg cells in lung tissue undergoing sterile damage are clonally expanded, and whether this relates to their tissue reparative functions, is currently unexplored.

The capacity to answer these questions regarding reparative Treg cells has been hindered by the inability to isolate live, repair-oriented Treg cells in a laboratory environment for downstream analyses. To address this deficiency, we generated a novel mouse strain that identifies Areg-producing Treg cells. Subsequently, we used this reporter strain to isolate reparative Treg cells in models of lung disease/damage for RNA and TCR sequencing. We utilized these datasets to identify previously unappreciated pathways relevant to Treg cell repair functions, characterize diseased-induced subsets of lung tissue Treg cells, and explore clonal expansion of Treg cells in response to sterile lung damage. Finally, we applied the findings from our sequencing studies to an in vitro assay of Treg cell–mediated repair, uncovering 4-1BB as a novel inducer of reparative activity.

## Results

### The Areg^Thy1.1^ reporter mouse delineates active Areg production from Treg cells during models of lung damage

Due to the inability to stain for Areg on the cell surface, isolation of live Areg-producing Treg cells has not been achievable. To create a way to isolate these cells, we created a novel murine reporter system: the *Areg^Thy1.1^* knock-in mouse. This was done by insertion of a sequence encoding a self-cleaving *Thy1^a^*(*Thy1.1*) construct into the endogenous *Areg* locus, wherein whenever a transcript of *Areg* is translated, *Thy1.1* is separately translated and trafficked to the surface of the cell for targeting by fluorescently conjugated antibodies (**Fig. 1A**, see Methods). We confirmed that these mice grow normally and maintain comparable histological features across multiple organ systems (**Fig. S1A-B**). Treg cells from spleens of *Areg^Thy1.1^* mice, after undergoing a short-term stimulation protocol, show a strong pattern of double-positive staining when stained for both Thy1.1 and endogenous AREG (**Fig. 1B**). To enable the isolation of live Treg cells, this strain was then crossed with *Foxp3^GFP^* mice to create *Foxp3^GFP^Areg^Thy1.1^*mice. Importantly, upon sorting and subjecting Treg cells to a long-term cytokine/TCR stimulation protocol, we saw no differences in proliferation or AREG production between Treg cells from *Foxp3^GFP^* mice and *Foxp3^GFP^Areg^Thy1^*^.*1*^ mice, demonstrating normal functionality of Treg cells harboring the *Areg^Thy1.1^* allele. (**Fig. S1C**).

**Figure 1.**
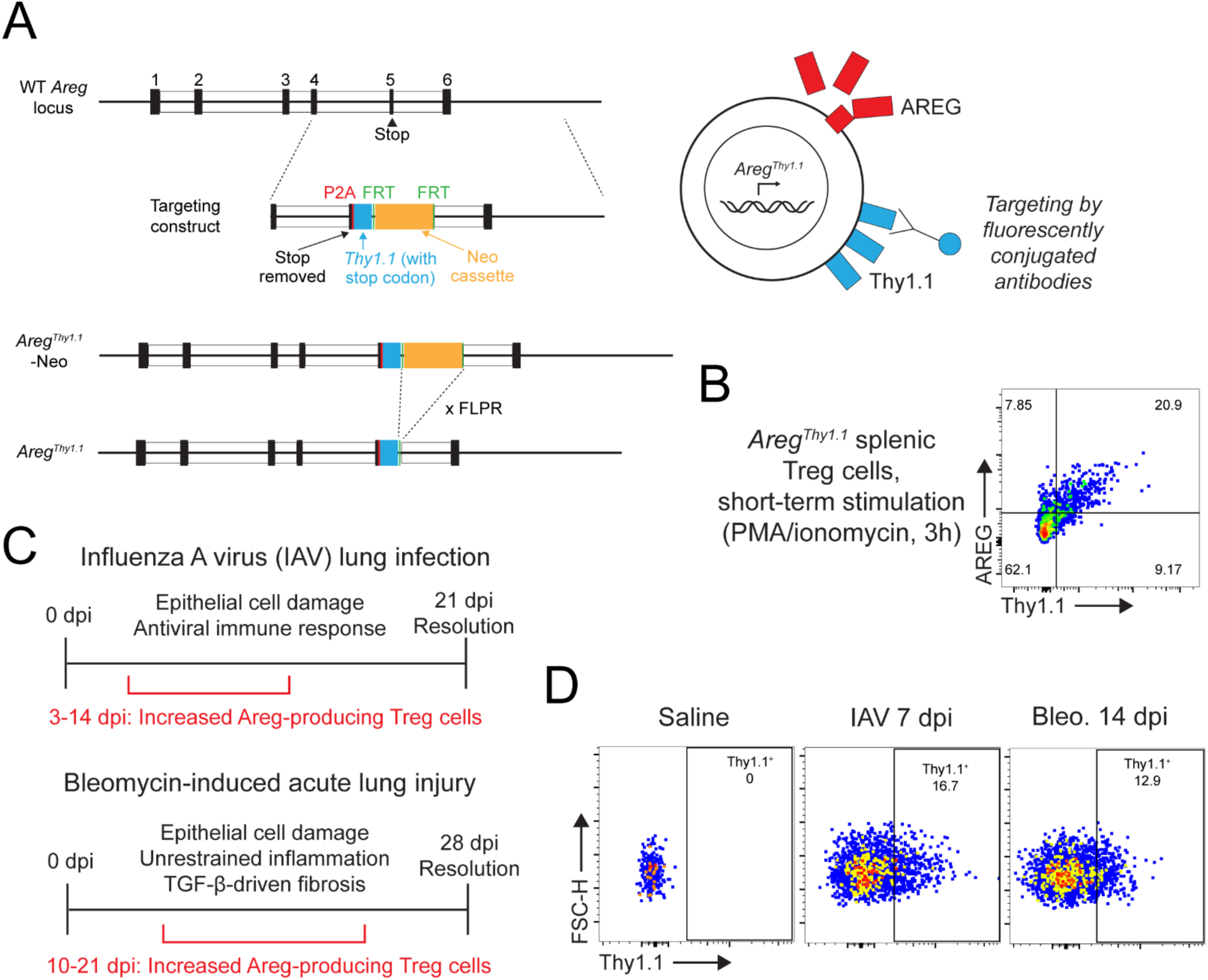
The *Areg^Thy1.1^* reporter mouse delineates active Areg production from Treg cells during models of lung damage. **A)** Left: schematic depicting genetic targeting of the endogenous *Areg* locus via homologous recombination with the P2A-*Thy1.1*-STOP-Neomycin knock-in construct, with subsequent crossing to the FLPR mouse to remove the neomycin cassette and create the final *Areg^Thy1.1^* allele. Right: A depiction of Areg transcription/translation in the *Areg^Thy1.1^* mouse; for each molecule of *Areg* mRNA translated, a single *Thy1.1* mRNA is also translated (as a separate protein, due to the P2A site) and traffics separately to the surface of the cell, where it can be targeted by fluorescently conjugated antibodies. **B)** Mouse splenocytes from *Areg^Thy1.1^* mice underwent a short-term stimulation protocol (PMA/ionomycin, 3h), then were stained for AREG using traditional methods (i.e., intracellularly, with biotinylated polyclonal antibody staining followed by streptavidin-mediated amplification) or for Thy1.1 (live staining with a fluorophore-conjugated antibody). Percentages of total Treg cells are shown in quadrants on plot. Representative staining shown from 2 experiments. **C)** Schematics depicting the models of lung damage used in this study, including general timecourse and disease characteristics, and Treg cell increases/Areg production status. **D)** Thy1.1 staining by flow cytometry on lung Tregs from *Areg^Thy1.1^* mice during the IAV or bleomycin (bleo.) models of lung damage, or from control (saline-treated) lungs. Days post-instillation (dpi) for each model indicated in figure. Representative staining from 2-3 experiments. See Fig. S2A for gating strategy throughout figure.

Areg production by lung Treg cells during influenza A virus (IAV) infection in mice has been previously shown by our group to be critical for proper tissue repair via signaling to a mesenchymal cell intermediate [6, 12]. The bleomycin model of sterile lung injury also induces high levels of Areg-producing Treg cells, as observed in several scRNA-seq datasets [13–15] and confirmed by our group at the protein level (data not shown). These models of lung pathology both involve extensive alveolar damage, a protracted immune response, and expansion of Treg cells. However, they differ in important and complementary ways with regards to method of injury, immunostimulatory antigens, and fibrosis induction [12, 16]. We thus focused our experiments on these models (**Fig. 1C**) and validated that *Foxp3^GFP^Areg^Thy1.1^*mice exhibited similar disease kinetics for both the IAV and bleomycin models compared to *Foxp3^GFP^* animals, as quantified by weight loss and body temperature (IAV and bleomycin), blood oxygen saturation (IAV), and lung Pdgfra^+^ mesenchymal cell induction of α-smooth muscle actin (αSMA) as a fibrosis indicator (bleomycin) (**Fig. S1D-E**).

In lung Treg cells from *Areg^Thy1.1^* mice treated with either IAV or bleomycin, we found a substantial increase in staining for Thy1.1 when compared to control saline-treated mice at 7 days post-instillation (dpi) for IAV and 14 dpi for bleomycin (**Fig. 1D** and **Fig. S2**). We additionally analyzed other cell types for Thy1.1 production during these models, finding negligible levels of induction for endothelial cells, mesenchymal cells, and myeloid cells (data not shown). Among lymphoid cells besides Treg cells, innate lymphoid cells (ILC) – previously shown to produce Areg [17] – display extensive Thy1.1 expression (**Fig. S1F** and **Fig. S2**); no other lymphoid cell types show appreciable expression (data not shown). Additionally, consistent with prior reports that epithelial cells are a major tissue source of Areg [18], lung epithelial cells (LEC) show extensive expression of Thy1.1 (**Fig. S1F** and **Fig. S2**). These characterizations of our new reporter mouse confirm that it provides an accurate and sensitive way to detect Areg-producing Treg cells and other cell types in live sortable cells.

### Bulk RNA-seq analysis of Areg-producing and non-producing lung Treg cells from IAV-or bleomycin-treated mice

We utilized our newly developed reporter mouse to sort live Thy1.1^−^ (Areg-producing) and Thy1.1^+^ (Areg-non-producing) Treg cells from the IAV and bleomycin models and performed bulk RNA-seq (**Fig. 2A).** For these models, timepoints of 8 dpi for IAV and 12 dpi for bleomycin were chosen as they represent the relative peaks of Treg expansion/Areg expression. Differential expression analysis between Thy1.1^−^ and Thy1.1^+^ Treg cells undergoing the IAV model exhibited 1,634 total differentially expressed genes (DEGs) using paired analysis (**Fig. 2B** and **Table S1**). For the bleomycin model, the same comparison exhibited 2,305 total DEGs. (**Fig. 2B** and **Table S2**). Confirming the efficacy of our reporter, *Areg* was a top DEG in each dataset.

**Figure 2.**
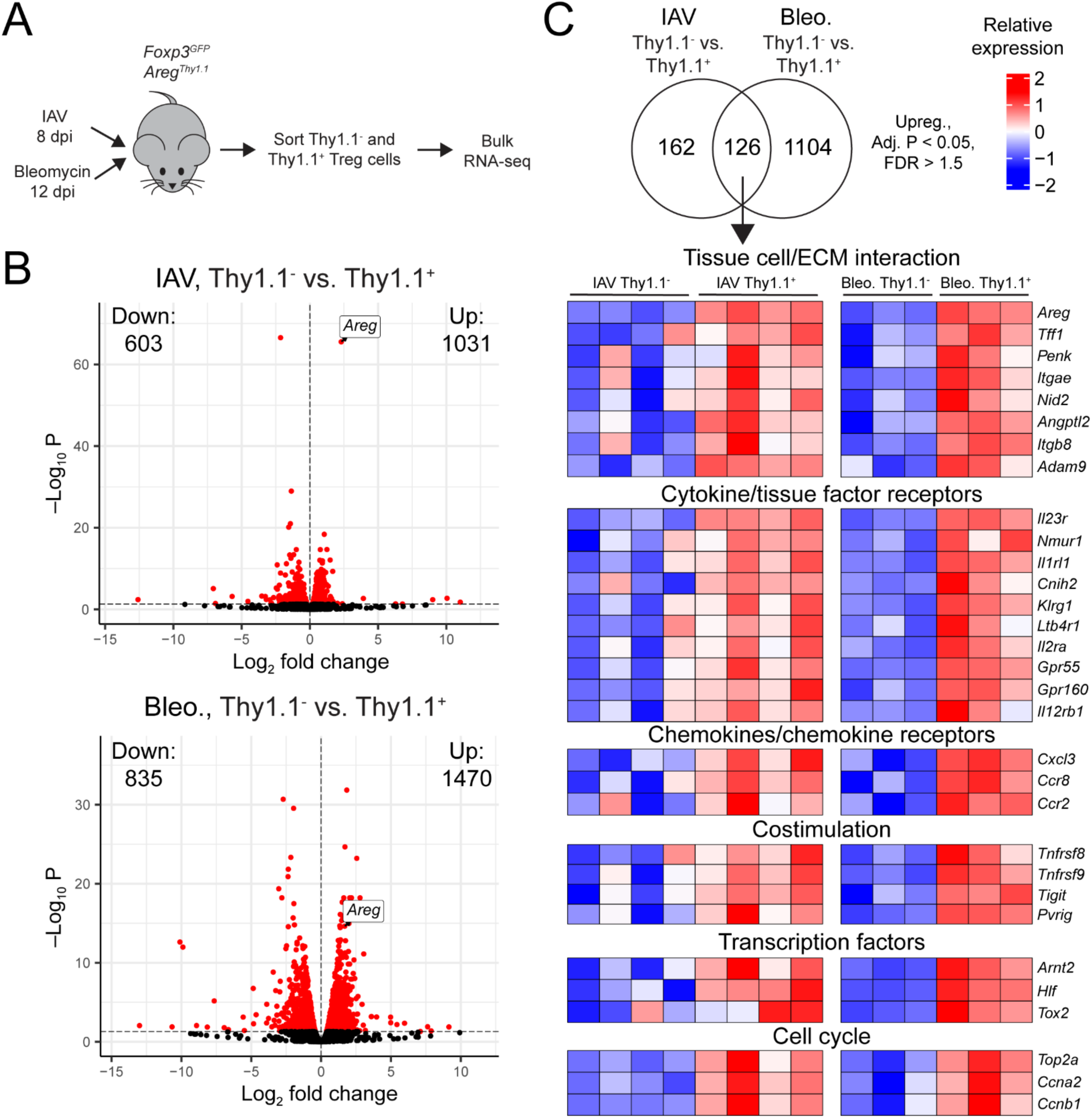
Bulk RNA-seq of Areg-producing and non-producing lung Treg cells from IAV-or bleomycin-treated mice. **A)** Schematic of bulk RNA-seq experiments using Thy1.1^−^ vs. Thy1.1^+^ lung Treg cells from IAV and bleomycin models. **B)** Volcano plots of differentially expressed genes (DEGs) for Thy1.1^−^ vs. Thy1.1^+^ lung Treg cells from bulk RNA-seq, from IAV and bleomycin models. Red dots on volcano plots: significant DEGs (FDR adj. p-value < 0.05). No fold change cutoff. Numbers of significantly upregulated and downregulated genes indicated on plots. **C)** Top: Venn diagram showing shared genes between IAV and bleomycin model comparisons that are significantly upregulated with a fold change induction of ≥1.5. Bottom: Heatmap of select shared genes, sorted by gene category; corresponding samples in each group are paired samples of Thy1.1^−^ vs. Thy1.1^+^ Treg cells from the same mouse.

At a more stringent fold change cutoff (FC > 1.5), we found that 126 upregulated DEGs were common between Thy1.1^−^ and Thy1.1^+^ Treg cells in the IAV and bleomycin models (**Fig. 2C**). This shared gene signature, by virtue of its commonality across both models, is most likely to directly relate to Treg cell activities associated with reparative cellular processes. Focusing on certain types of genes within this signature, we found that several genes for tissue cell/ECM interaction mediators, receptors for various cytokines and tissue factors, and chemokines/chemokine receptors were significantly altered across both models. Significantly increased presence of transcription factors *Arnt2*, *Hlf*, and *Tox2* was also seen across models, indicating an altered gene regulatory state of Areg-producing cells. Interestingly, several genes for costimulatory molecules (*Tnfrsf8* [CD30], *Tnfrsf9* [4-1BB], *Tigit*, and *Pvrig*) were commonly differentially expressed in both models; costimulation of Treg cells has not been previously identified as a mediator of tissue repair functionality. We additionally saw consistent upregulation of several cell cycle genes in Thy1.1^+^ Treg cells; however, this most likely reflects heterogeneity in sorted Thy1.1^−^ and Thy1.1^+^ Treg cells, wherein more proliferating cells within the latter group bias this analysis.

### scRNA-seq analysis of Areg-producing and non-producing lung Treg cells from IAV-or bleomycin-treated mice

To account for heterogeneity within the Treg cell population, we performed scRNA-seq on sorted Thy1.1^−^ and Thy1.1^+^ Treg cells from the lungs of mice undergoing the IAV and bleomycin models, as well as Treg cells from control mice (saline-treated) (all Thy1.1^−^) (**Fig. 3A**). We chose a different timepoint from the bulk RNA-seq dataset for the IAV model (5 dpi), since this has previously been shown to be the timepoint when Treg cells are the dominant source of Areg in lung tissue [6]; the same 12 dpi timepoint was used for the bleomycin model as in the bulk RNA-seq dataset. Cells from separate mice were hashed to permit identification of cells on a per-mouse basis for analyses. Notably, Thy1.1^−^ and Thy1.1^+^ cells were run in parallel on separate lanes, in order for us to be able to identify these cells without a separate round of cell hashing. In doing so, we included roughly similar amounts of Thy1.1^−^ and Thy1.1^+^ cells to give us the ability to better determine heterogeneity and representation within the Thy1.1^+^ group. However, the consequence of this strategy is that cell counts from the groups in these experiments are not reflective of the actual tissue proportions of Treg cells from different subtypes; thus, we did not attempt to perform analyses of amounts of Treg cells of each subtype from these datasets.

**Figure 3.**
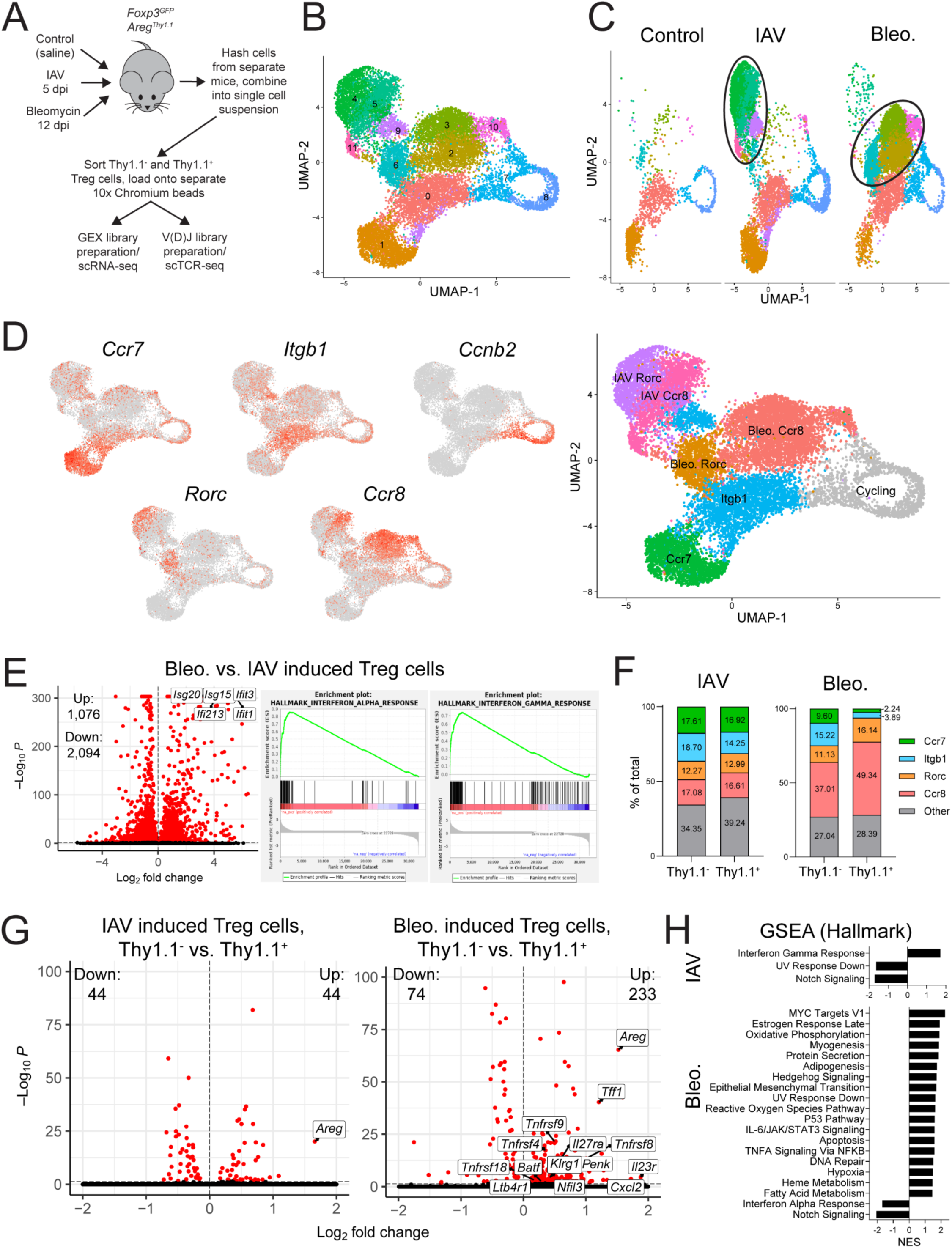
scRNA-seq of Areg-producing and non-producing lung Treg cells from IAV-or bleomycin-treated mice. **A)** Schematic of single cell (sc) RNA-seq experiments using Thy1.1^−^ vs. Thy1.1^+^ lung Treg cells from IAV and bleomycin models (as well as control, saline-treated mice). **B)** UMAP of clustered cells from sc gene expression analysis using Seurat. **C)** UMAP from (B), split by treatment status of Treg cells. Circles highlight groups that are largely specific to each type of tissue damage model (IAV or bleomycin). **D)** Left: Feature plots of several genes uncovered by marker gene analysis of groups from (B). Using these genes and/or by presence in a specific model (IAV or bleomycin), identities were assigned to groups of cells (re-assigned UMAP on right). **E)** Left: Volcano plot of DEGs between induced Treg cell subgroups from the IAV model (“IAV Rorc” and “IAV Ccr8” combined) vs. from the bleomycin model (“Bleo. Rorc” and “Bleo. Ccr8” combined). Red dots: significantly differentially expressed (FDR adj. p-value < 0.05). No fold change cutoff. Numbers of significantly upregulated and downregulated genes indicated on plots. Right: GSEA enrichment plots of top 2 significant pathways in the gene signature shown here, using Hallmark curated gene sets. Both pathways significant at FDR q-value < 0.05. **F**) Proportions of each assigned subgroup in Thy1.1^−^ vs. Thy1.1^+^ Treg cells (see Fig. S2E for individual reclustering from each model, used to assign cells to subgroups in these plots). **G)** Volcano plots of DEGs from Thy1.1^−^ vs. Thy1.1^+^ induced Treg cells (Ccr8 and Rorc subgroups combined) from IAV and bleomycin models. Red dots on volcano plots: significant DEGs (FDR adj. p-value < 0.05). No fold change cutoff. Numbers of significantly upregulated and downregulated genes indicated on plots. **H)** Pathway analysis using GSEA, in full gene signatures from IAV or bleomycin datasets depicted in (B) (from Hallmark curated gene sets). NES: Normalized enrichment scores. All values displayed are significant at FDR q-value < 0.05.

Upon initial analysis using the Seurat platform [19], we first noted that most of the cells show high expression levels of *Foxp3* and cluster in the same area of UMAP space, confirming their identity as Treg cells, with a much smaller subcluster showing markers for other cell types (epithelial cells [*Sftpc*], endothelial cells [*Cldn5*], macrophages [*Chil3*]) (**Fig. S3A**); this contaminating subcluster was excluded from further analyses. To confirm that our *Areg^Thy1.1^* reporter mouse was effective in allowing us to preferentially isolate Areg-producing Treg cells, we queried *Areg* gene expression in total Thy1.1^−^ and Thy1.1^+^ Treg cells and found that it was highly increased on a per-mouse basis, again validating the efficacy of our reporter (**Fig. S3B**). Upon re-clustering with contaminating cells excluded, we found that Treg cells were partitioned into 11 subclusters (**Fig. 3B**). Strikingly, when separating out control, IAV-infected, and bleomycin-treated Treg cells, we found that certain clusters were present across all conditions, while other clusters appear to be largely specific to IAV infection or bleomycin induction (circled in **Fig. 3C**).

We then assessed each population for various marker genes, assigning cell populations based on certain genes with specific expression as well as the damage model of origin (**Fig. 3D**). The clusters marked by *Ccr7*, *Itgb1*, and *Ccnb2* are present across all conditions, including controls. *Ccr7* expression has previously been indicated as a marker for naïve Treg cells [20]. While mouse lungs used to isolate Treg cells underwent perfusion prior to lung processing and sorting, flow cytometry experiments with mouse lungs prepared in the same way, with intravenous (i.v.) labelling to mark circulating cells, indicated that some blood Treg cells remain in lung tissue even after perfusion (**Fig. S3C**). Thus, we concluded that our “Ccr7” subset is most likely circulating naïve Treg cells. Since the *Itgb1*^+^ Treg cell population is present in control mice in addition to mice undergoing damage models, and is located closest to *Ccr7*^+^ circulating Treg cells in UMAP space, we chose to interpret the “Itgb1” group as the baseline tissue-adapted lung Treg cell population seen in previous datasets of lung Treg cells from healthy mice [21]. *Ccnb2*^+^ Treg cells demonstrated high expression of cell cycle genes (termed “Cycling” in our group labels) and were therefore excluded from further analysis. One of the groups that only appears in the settings of tissue injury is marked by the *Rorc* gene, encoding transcription factor RORγt associated with Th17 cells and Treg cells of the gastrointestinal (GI) tract [22]. A recent report indicates that RORγt^+^ Treg cells can migrate from the GI tract to the lungs under certain conditions [23], although other reports have indicated that they can be derived peripherally at locations other than the GI tract [24]. The other damage-induced group is marked by the gene *Ccr8*, which has been previously described as a marker for tissue-adapted, Th2-like Treg cells [25]; Treg cells with a similar signature have been repeatedly described in lung tumors and other types of cancer [26, 27].

We looked at the expression of several known Treg cell– and T cell–related genes to gain a better understanding of the nature of these subgroups (**Fig. S3D**). *S1p1r*, the receptor which allows naïve T cells to follow cues for recruitment to tissue sites, is highest in the Ccr7 subgroup, confirming their identity as naïve, likely circulating Treg cells. Classical Treg cell immunosuppression mediators *Il10*, *Tgfb1*, and *Ctla4* show generally higher expression in tissue-resident subsets (Itgb1, Rorc, and Ccr8), implying that these groups may simultaneously perform immunosuppression and tissue repair functions. We further assessed Treg cell subgroup expression of master T helper (Th) cell transcription factors *Tbx21* (T-bet, characteristic of Type 1/Th1 immunity), *Gata3* (characteristic of Type 2/Th2 immunity), and *Rorc* (ROR-γt, characteristic of Type 3/Th17 immunity) [28]. We found that *Tbx21* was not strongly expressed in any subgroup, while Ccr8 and Rorc subgroups had strongest expression of *Gata3* and *Rorc*, respectively. Finally, *Ikzf2* (encoding transcription factor Helios) has been proposed to be elevated in thymic-derived Treg cells in comparison to peripherally-derived Treg cells [29]. We see heightened *Ikzf2* expression on Itgb1 and Ccr8 subgroups, with comparably lower expression on Rorc subgroups in each model, suggesting that the Itgb1 and Ccr8 subgroups may be thymically-derived, whereas Rorc subgroups may be peripherally-induced.

Notably, the IAV– and bleomycin-induced Treg cells (i.e., not present in control mice) (**Fig. 3C**) each consist of two main clusters marked by similar genes (*Ccr8* and *Rorc*) (**Fig. 3D**). To ascertain why these IAV– and bleomycin-specific cells did not cluster in similar UMAP space despite this seemingly analogous gene expression pattern, we directly compared gene expression in bleomycin-induced vs. IAV-induced Treg cells (Ccr8 and Rorc clusters combined) (**Fig. 3E** and **Table S3**). We found that the primary axis of difference between these cells is the induction of interferon inducible genes, which may be expected from Treg cells in the interferon-enriched environment of IAV infection. However, the similar bifurcation of IAV– and bleomycin-induced Treg cells into Ccr8 and Rorc subsets indicates that in either model, newly emergent lung Treg cells are likely of similar subtypes.

To gain greater clustering resolution on induced Treg cell subsets, we performed separate clustering on cells only from the IAV or bleomycin models (**Fig. S3E**). This allowed greater specificity in defining the primary Treg subgroups (Ccr7, Itgb1, Rorc, and Ccr8) seen in each model. Additionally, there were cells that showed a lack of markers for our specific subgroups, which were labelled as “Undefined/Intermediate” populations (“Undef./Int.”) and excluded from future analyses. Interestingly, when we performed separate clustering on Treg cells from control mice (**Fig. S3E**), beyond the Ccr7, Itgb1, and Cycling groups visible in **Fig. 3C**, we also detected small groups of *Rorc*– and *Ccr8*-expressing cells.

To determine if the proportions of subtypes of Treg cells seen in this analysis are related to their Areg production status, we analyzed the subgroup composition of Thy1.1^−^ and Thy1.1^+^ cells (**Fig. 3F**). In the IAV model, there were only minor changes in the composition of each subset of Treg cells (no changes > 5%). However, in the bleomycin group, we found that there were substantial increases in the Rorc and Ccr8 subgroups in Thy1.1^+^ Treg cells compared to Thy1.1^−^, with corresponding reductions in the Ccr7 and Itgb1 subgroups.

Referring back to a major potential advantage compared with the bulk RNA-seq studies we conducted, the subclustering of baseline, proliferating, and damage-induced Treg cells from these datasets allowed us to strictly isolate damage-induced cells for comparison between Thy1.1^−^ and Thy1.1^+^ populations (unlike in the bulk RNA-seq dataset, where contamination with proliferating cells seemed to alter our DEG signatures). With this in mind, we performed DEG analysis on Thy1.1^−^ and Thy1.1^+^ cells from induced Treg cell subsets (Ccr8 and Rorc clusters combined), from both the IAV and bleomycin datasets (**Fig. 3G, Table S4**, and **Table S5**). We found that both models exhibited altered gene signatures between these cells, with a higher level of transcriptional alteration in the bleomycin comparison vs. the IAV comparison. The bleomycin gene signature includes several genes identified previously in our bulk RNA-seq dataset, including known Treg cell repair mediators (*Areg*, *Tff1*, *Penk*) [9, 30], tissue factor receptors (*Klrg1*, *Il23r*, *Ltb4r1*), and costimulation mediators (*Tnfrsf8*, *Tnfrsf9*); other genes in these classes were novel to this more precise analysis (i.e., not represented in bulk analyses) (*Il27ra*, *Tnfrsf4*, *Tnfrsf18*). Additionally, chemokine *Cxcl2* is strongly induced in this context, pointing to a potential role for reparative Treg cells in recruiting a novel immune milieu to the site of tissue injury. Transcription factors *Batf* and *Nfil3*, previously identified as important for tissue-adapted identity in Treg cells [31, 32], were also upregulated.

Pathway analysis of these gene signatures revealed only 3 significantly altered pathways in the IAV scenario, while there were 20 significantly altered pathways in the bleomycin comparison (**Fig. 3H**). Among the novel pathways revealed by this approach, several pathways (“Hedgehog Signaling”, “IL-6/JAK/STAT3 Signaling”, “Reactive Oxygen Species Pathway”, and “Fatty Acid Metabolism”) are novel modes of signaling ascribed to reparative Treg cells in this context. Interestingly, significant downregulation of the “Notch Signaling” pathway is apparent in both the IAV and bleomycin datasets; Notch signaling has recently been shown to be antagonistic towards lung Treg cell repair activity/Areg production [33].

As discussed in the next section, we found that both our IAV and bleomycin Treg cells from this analysis showed some degree of clonal expansion, but not enough for substantial analysis of expanded Treg cell clones. Thus, we added an additional dataset to our investigations – analysis of bleomycin-induced lung Treg cells at 21 dpi (**Fig. S4A**), when we anticipated that Treg cell clonal expansion would be more pronounced. Gene expression analyses gleaned from scRNA-seq of this new dataset yielded similar results to those shown for the bleomycin 12 dpi dataset, with similar clustering of Treg cell subsets (**Fig. S4B-D**), similar overrepresentation of Thy1.1^+^ cells in induced Treg cell subgroups (**Fig. S4E**), and similar gene expression differences between Thy1.1^−^ and Thy1.1^+^ cells in the induced Treg cell subgroups (**Fig. S4F** and **Table S6**). In comparison to the bleomycin 12 dpi dataset, significant DEGs at this later timepoint (when Treg cells have been maintained in tissue for a longer duration) include similar transcription factors to those seen in our bulk RNA-seq (*Arnt2*, *Tox2*), further validating these in the induction of a tissue-reparative program in Treg cells. There is also a significant upregulation of *Cd44*, which may implicate the establishment of a memory-like reparative Treg cell population at this later timepoint.

### scTCR-seq of Areg-producing and non-producing lung Treg cells from IAV-or bleomycin-treated mice

We simultaneously evaluated the TCR repertoire of tissue Treg cells during these disease models at single cell resolution. Throughout this section, consistent with prior studies [34–36], we perform comparisons using CDR3β amino acid sequence data; while information regarding CDR3α status, nucleotide sequences, and V/D/J gene usage are available from our data, since CDR3β have heightened potential for diversification due to potential inclusion of the D region/more sites for nucleotide addition [37], and since CDR3β chains form the dominant contact with epitope in most TCRs [38], evaluation of only CDR3β sequences provided sufficient analytic depth. Thus, for the remainder of this section, “TCR” refers to unique CDR3β amino acid sequences.

On a per-mouse basis, TCR diversity of Treg cells in each dataset was highest in control mice, while a substantial decrease was seen in IAV 5 dpi mice, with an even further decrease seen in bleomycin-treated mice (at both 12 dpi and 21 dpi). (**Fig. 4A**). Next, we evaluated Treg clonal expansion between each separate model/timepoint (**Fig. 4B**). As expected, control mice showed minimal clonal expansion with no clones expanded to >10 cells. Treg cells from IAV treated mice (5 dpi) also showed minimal expansion with only a small amount of clones expanded >10, likely as a consequence of the early nature of this timepoint. Within the bleomycin datasets, we see several clones expanded to >10 at the 12 dpi timepoint, while this is even further pronounced at the 21 dpi timepoint (with several clones expanded to >100). We additionally evaluated TCR expansion in Thy1.1^−^ and Thy1.1^+^ cells from each separate model/timepoint (**Fig. 4C**). While there were slight increases in expansion of clones in Thy1.1^+^ Treg cells compared to Thy1.1^−^ in the IAV 5 dpi and bleomycin 12 dpi datasets, this difference is profoundly increased in the bleomycin 21 dpi dataset, where the majority of Thy1.1^+^ cells are expanded. These data indicate that Treg cells undergo progressive clonal expansion in a sterile model of lung damage (bleomycin), and that Areg-producing reparative Treg cells show features of heightened clonal expansion.

**Figure 4.**
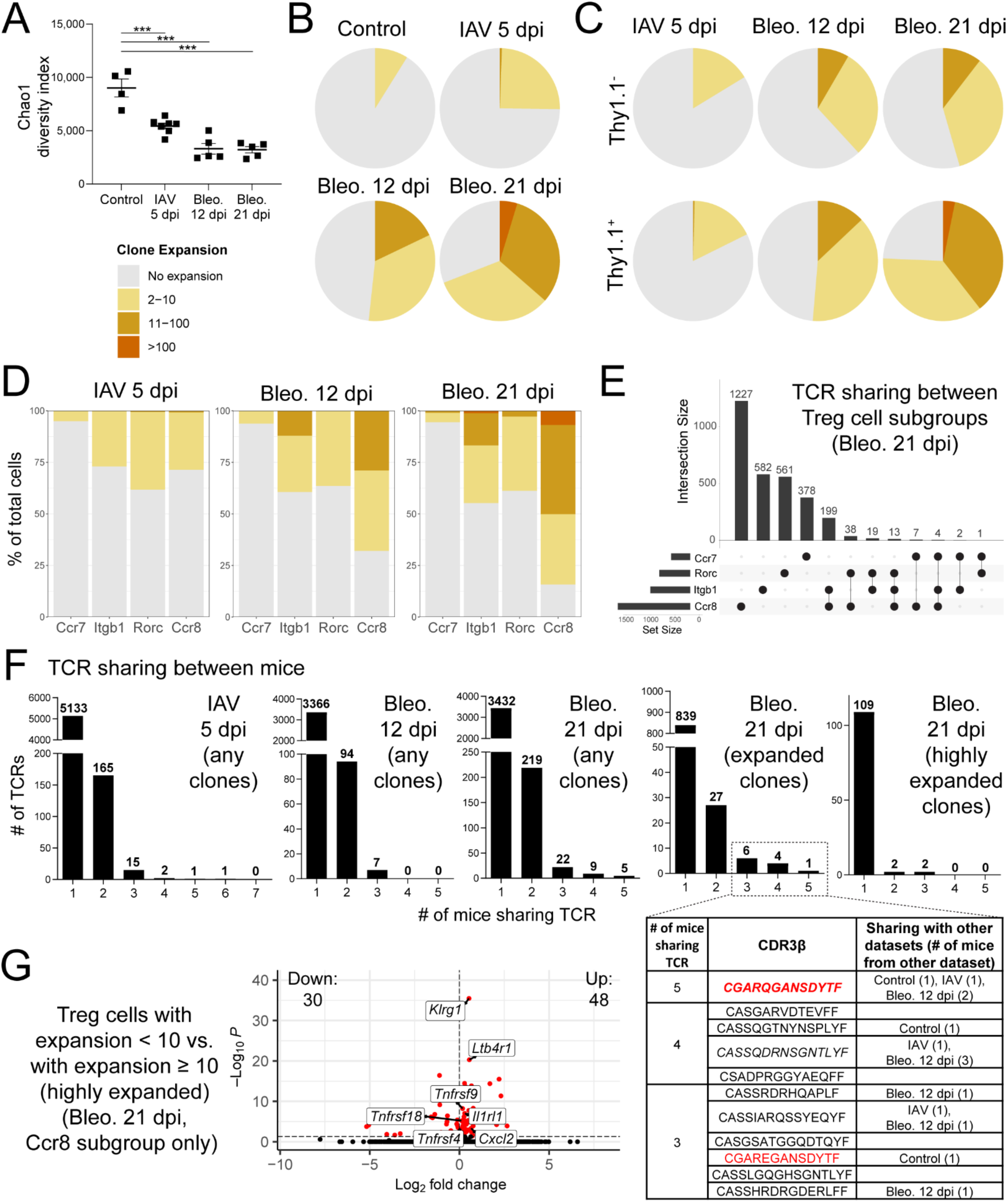
scTCR-seq of Areg-producing and non-producing lung Treg cells from IAV-or bleomycin-treated mice. **A)** Chao1 diversity scores for the full Treg cell TCR repertoire (by CDR3β) of separate mice in each treatment dataset. **B)** Pie charts representing the clonal expansion status of all Treg cells with detected CDR3β sequences within each treatment dataset. **C)** Pie charts representing the clonal expansion status of all Treg cells with detected CDR3β sequences within each treatment dataset, broken down by Thy1.1^−^ vs. Thy1.1^+^ status of Treg cells (control mice were not included, due to their being all Thy1.1^−^). **D)** Stacked bar plots representing the clonal expansion status of Treg cells with detected TCR CDR3β sequences within each treatment dataset, subdivided by subgroup of Treg cell determined from our previous clustering. Only the 4 main subgroups of Treg cells were included (Ccr7, Itgb1, Rorc, Ccr8). **E)** Upset plot depicting sharing of TCR CDR3β sequences between subgroups of Treg cells in the bleomycin 21 dpi dataset. Connections between dots indicate sharing between subgroups. **F)** Sharing of TCR CDR3β sequences between mice in each treatment group. Numbers over bars indicate total number of TCRs that are shared by n mice (indicated by the x-axis). Whether this sharing considers any clones (including unexpanded, singlicate clones), expanded clones (≥2 clones in each mouse), or highly expanded clones (≥10 clones in each mouse) is indicated in the top right of each plot. Inset/table: specific CDR3β sequences that are expanded (≥2 clones in each mouse) in at least 3 mice from the bleomycin 21 dpi dataset, and whether they are shared with any clones from the other treatment datasets. Red: TCR clones investigated in Rappazzo 2020, see Results section. Italicized: TCR clones identified in silica-treated mouse lungs in Bao et al. 2022, see Results section. **G)** Volcano plot of DEGs between highly expanded Treg cells (≥10 clones in full dataset) vs. less expanded Treg cells (<10 expanded clones), in the bleomycin 21 dpi Ccr8 subgroup. Red dots: significantly differentially expressed (FDR adj. p-value < 0.05). No fold change cutoff. Numbers of significantly upregulated and downregulated genes indicated on plots. Standard error displayed on graphs; n.s: not significant, *: 0.01<p<0.05, **: 0.001<p<0.01, ***: 0.0001<p<0.001, ****: p<0.0001.

Simultaneous evaluation of scRNA-seq and scTCR-seq additionally allowed us to examine TCR expansion in specific Treg cell subsets. These results indicate that the Ccr8 subset in the bleomycin model (at either timepoint) contains the greatest amount of expanded Treg cells, while there do not appear to be major differences among Itgb1, Rorc, and Ccr8 subsets in the IAV dataset (**Fig. 4D**). With respect to clonal sharing between subgroups, we found that the highly expanded Ccr8 TCRs in the bleomycin 21 dpi dataset primarily are shared with the Itgb1 subgroup, with lesser sharing in other subgroups (**Fig. 4E**). This may indicate a developmental trajectory between these subgroups, wherein Treg cells first entering tissue are of the Itgb1 phenotype, later converting to a Ccr8 phenotype after further clonal expansion; this interpretation is supported by the similar expression of *Gata3* and *Ikzf2* between the Itgb1 and Ccr8 subgroups (compared to Rorc) (**Fig. S3D**).

Next, we sought to assess TCR sharing between different mice undergoing each damage model/timepoint (**Fig. 4F**). For the IAV and the bleomycin 12 dpi dataset, no TCRs were shared among all mice analyzed, although there was a lower degree of sharing between 3–6 mice in each model. However, for the bleomycin 21 dpi dataset, 5 TCRs were shared among all mice, with an additional 31 shared in at least 3 mice. We then heightened the stringency of this analysis to ascertain the prevalence of expanded TCRs (≥2 clones/mouse) across mice in the bleomycin 21 dpi dataset. When looking at only expanded TCRs, we found that 11 were shared by at least 3 mice from this dataset (highlighted in inset/table in **Fig. 4F**). Notably, several of these clones were found in control, IAV 5 dpi, and bleomycin 12 dpi datasets. To query whether this TCR sharing pattern is apparent among TCRs with the highest degree of clonal expansion, we then performed this same analysis on highly expanded clones from the bleomycin 21 dpi dataset (≥10 clones/mouse) and found that none were shared by all or even 4/5 mice; only 2 were shared by 3 mice. This latter analysis seems to imply that the expansion of highly expanded clones is to some degree stochastic, with different highly expanded clones dominating in different mice. On the other hand, TCR diversity generation in vivo has incredible breadth, and we have only analyzed a limited amount of Treg cells from each mouse in this study. In fact, in order to perform analyses of Treg cell repertoires, previous studies have used murine genetic systems to artificially limit T cell diversity [39, 40]. Therefore, the finding that several TCRs are shared at the “expanded” level (inset/table in **Fig. 4F**) may instead be taken as evidence that commonly expanded clones are indeed present in response to tissue damage. To this effect, 2 of the identified shared/expanded clones (italicized in table) were identified as highly expanded T cell clones in a model of silica-induced lung fibrosis [36], and 2 (red in table) were found to be highly expanded Treg cell clones in a model of immunogenic lung cancer [41]. This finding suggests that certain common clones in our dataset may arise in response to lung damage in general, rather than to an aspect specific to the model of bleomycin-induced ALI.

Lastly, we used the high prevalence of expanded clones in our bleomycin 21 dpi dataset to evaluate potential gene signature differences between highly expanded Treg clones (≥10 clones across all cells/mice) and other Treg cells in this damage context (**Fig. 4G** and **Table S7**). This comparison yielded a DEG signature similar to that seen for our comparisons between Areg-producing and –non-producing Treg cells, with upregulation in receptors for tissue factors (*Klrg1*, *Il1rl1*, *Ltb4r1*), costimulation mediators (*Tnfrsf4*, *Tnfrsf9*, *Tnfrsf18*), and *Cxcl2*.

### Analysis of immunosuppression and tissue repair gene modules and functional activity of Areg-producing and non-producing lung Treg cells from IAV-or bleomycin-treated mice

A major question since the discovery of the reparative function of Treg cells has been whether distinct subsets of Treg cells are responsible for immunosuppression vs. tissue repair functionality. We attempted to evaluate this paradigm by querying our scRNA-seq datasets using gene module analysis. We generated an “Immunosuppression” gene module that includes genes known to relate specifically to immunosuppressive functions in Treg cells (**Fig. 5A**) [1, 42]. We also created a “Tissue repair” gene module that includes genes that have been shown in published literature to encode mediators produced by Treg cells that have a direct impact on tissue repair [2], as well as the transcription factor *Pparg*, which has been demonstrated to be a marker of tissue-adapted Treg cells [43] (**Fig. 5A**). We found that our gene modules for both “Immunosuppression” and “Tissue repair” were both primarily enriched in the Ccr8 subset in each of our 3 separate datasets (IAV 5 dpi, bleomycin 12 dpi, bleomycin 21 dpi) (data not shown). Thus, we chose to further focus on only the Ccr8 subset for these analyses.

**Figure 5.**
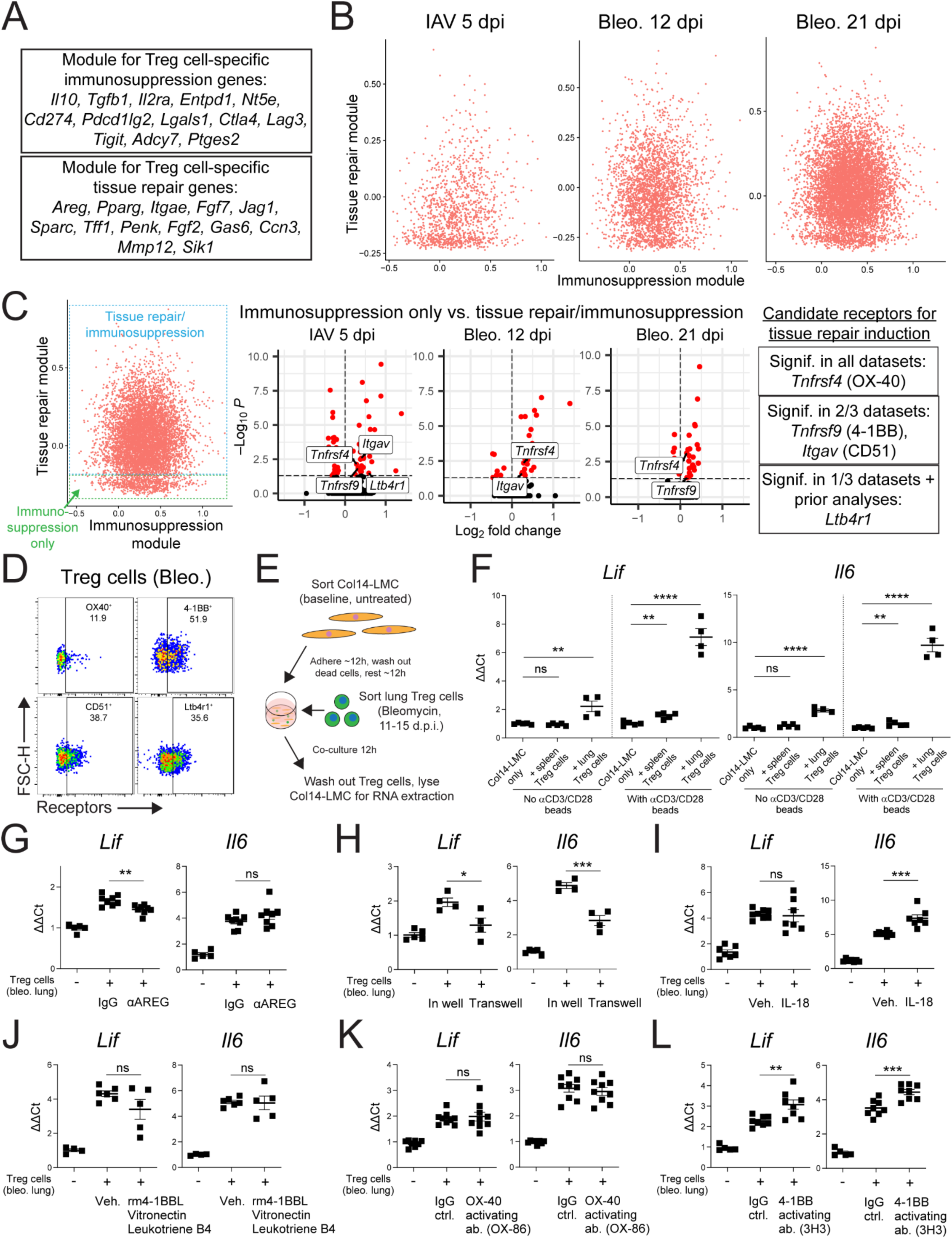
Analysis of immunosuppression and tissue repair gene modules and functional activity of Areg-producing and non-producing lung Treg cells from IAV-or bleomycin-treated mice. **A)** “Immunosuppression” and “Tissue repair” modules of genes (Treg-specific), established from literature. **B)** “Immunosuppression” and “Tissue repair” module scores were calculated for cells in the Ccr8 subgroup of each treatment dataset, then plotted against each other. **C)** Left: Subclusters with differing patterns of module expression are indicated by boxes on plot. Middle: Volcano plots of DEGs between groups indicated at left for each treatment dataset. Red dots: significantly differentially expressed (FDR adj. p-value < 0.05). No fold change cutoff. Genes encoding receptors of interest for activation of tissue repair function are indicated on graphs and in the summary on the right. **D)** Flow cytometric profiling of lung Treg cells from bleomycin-treated lung tissue (11-15 dpi), for receptors identified in (C). **E)** Experimental schematic of Col14-LMC and Treg cell co-culture. **F)** qPCR for Treg cell-induced genes *Lif* and *Il6* in Col14-LMC following lung or spleen Treg cell co-culture, with or without αCD3/CD28 beads for T cell activation as indicated. **G)** qPCR for *Lif* and *Il6* in Col14-LMC following Treg cell co-culture, with control IgG or αAREG antibody. **H)** qPCR for *Lif* and *Il6* in Col14-LMC following Treg cell direct co-culture, or separation with a 0.4 µm transwell insert. **I)** qPCR for *Lif* and *Il6* in Col14-LMC following Treg cell co-culture, with vehicle or rmIL-18. **J)** qPCR for *Lif* and *Il6* in Col14-LMC following Treg cell co-culture, with vehicle or a combination of rm4-1BB ligand, vitronectin, and leukotriene B4. **K)** qPCR for *Lif* and *Il6* in Col14-LMC following Treg cell co-culture, with control IgG or αOX-40 activating antibody (clone OX-86). **L)** qPCR for *Lif* and *Il6* in Col14-LMC following Treg cell co-culture, with control IgG or α4-1BB activating antibody (clone 3H3). Standard error displayed on graphs; n.s: not significant, *: 0.01<p<0.05, **: 0.001<p<0.01, ***: 0.0001<p<0.001, ****: p<0.0001.

Cellular scores from the “Immunosuppression” module and the “Tissue repair” module were plotted against each other within the Ccr8 subset for each dataset (**Fig. 5B**). Strikingly, we saw that two separate groups of cells appear: a group that expresses immunosuppression genes but no tissue repair genes (which we called the “Immunosuppression only” group), and a group that expresses both immunosuppression genes and tissue repair genes (which we entitled the “Tissue repair/immunosuppression” group) (**Fig. 5C**). To gather further insight into the differences between these subgroups, we performed differential expression analysis comparing them in each dataset (**Fig. 5C** and **Table S8-S10**). When analyzing these gene signatures, our focus was on discovering potential receptors on “Tissue repair/immunosuppression” Treg cells that can potentially activate their tissue repair functionality. To this effect, we found that *Tnfrsf4*, encoding T cell costimulatory molecule OX-40, was significantly upregulated in all three datasets; *Tnfrsf9* and *Itgav*, encoding T cell costimulatory molecule 4-1BB and CD51 (integrin αV, a receptor for extracellular matrix [ECM] molecule vitronectin), were significantly upregulated in 2/3 datasets; and *Ltb4r1*, encoding leukotriene B4 receptor, was significantly upregulated in 1/3 datasets (but was additionally differentially expressed in other datasets throughout this study). These receptors – reflective of costimulation by other cells in the lung milieu (OX-40 and 4-1BB), ECM-induced tissue adaptation (CD51), or leukotriene signaling (Ltb4r1) – have not been previously explored for their role in inducing tissue repair by Treg cells. We confirmed expression of these proteins in bleomycin-induced lung Treg cells (**Fig. 5D**) and found low levels of OX-40, with higher levels of 4-1BB, CD51, and Ltb4r1.

To address the potential for activation of tissue repair function of these receptors, we utilized an ex vivo co-culture assay previously developed in our lab by which to test the tissue repair functionality of Treg cells. Here, Treg cells are co-cultured with a sub-population of lung mesenchymal cells (LMC) involved in alveolar regeneration (characterized by high expression of the *Col14a1* gene encoding Collagen XIV; hereafter referred to as “Col14-LMC”) (**Fig. 5E** and **Fig. S2**) [12]. Co-culture of bleomycin-induced lung Treg cells promoted greater expression of *Il6* and *Lif* (another IL-6 family cytokine) from Col4-LMC, compared to Col14-LMC alone or splenic Treg cells from bleomycin-treated mice (**Fig. 5F**). This was further enhanced by the addition of αCD3/CD28 beads to cultures, in order to stimulate Treg cell activity (**Fig. 5F**). Blocking of Areg with an αAreg antibody significantly reduced the transcription of *Lif* (but not *Il6*) in Col14-LMC compared to IgG controls (**Fig. 5G**). The partial nature of this inhibition indicates that there are additional non-Areg mechanisms of Col14-LMC stimulation at play. We additionally tested contact dependency of Treg cell-mediated Col14-LMC activity by separating Treg cells from Col14-LMC in a transwell plate. We found that there was a partial but significant decrease in *Il6* and *Lif* transcription in transwell-separated Col14-LMC compared to controls (**Fig. 5H**). The partial nature of this decrease indicates that there are likely both contact-dependent and soluble factors from Treg cells that contribute to Col14-LMC stimulation.

To assess whether factors known to induce the tissue repair activity of Treg cells are able to stimulate Treg cell-mediated Col14-LMC activity, we tested the addition of IL-18 in this assay. IL-33 was not able to be tested, because it unexpectedly caused direct stimulation of Col14-LMC (data not shown). IL-18, when added along with bleomycin-induced lung Treg cells, was able to significantly increase transcription of *Il6* (but not *Lif*) in Col14-LMC compared to vehicle controls (**Fig. 5I**). Interestingly, this assay does not seem to be simply demonstrating non-specific effects of Treg cell activation (i.e., wherein other Treg cell functions are also amplified), since IL-18 has previously been demonstrated to dampen Treg cell-mediated suppression [44].

We then turned to testing the potential repair activity-inducing ligands for the receptors uncovered from our scRNA-seq analyses (**Fig. 5C**). We first attempted to treat Treg cells in the co-culture assay to ligands for 4-1BB (recombinant mouse 4-1BBL), CD51 (vitronectin), and Ltb4r1 (leukotriene B4), based on their substantial expression levels on bleomycin-induced lung Treg cells (**Fig. 5D**). The combination of these ligands was unable to confer greater Treg cell–induced Col14-LMC activity, based on unchanged expression of *Il6* and *Lif* (**Fig. 5J**). While OX-40 was expressed at a comparatively lower level than the other receptors at the protein level, it showed the most consistent upregulation in our scRNA-seq analysis of tissue repair/immunosuppression-oriented vs. immunosuppression only–oriented Treg cells, so we decided to test activity through this costimulator molecule as well, using an activating antibody towards OX-40. However, this approach also yielded no changes at the level of Col14-LMC transcription compared to IgG controls (**Fig. 5K**).

Maximal 4-1BB downstream activation requires multimerization via interaction with multiple 4-1BBL proteins on the surface of costimulating cells [45], an effect that may not be occurring when recombinant 4-1BBL is used in this system as in **Fig. 5J**. Thus, we used an activating antibody for 4-1BB that has previously shown efficacy towards T cell activation in vivo [46], to induce some level of multimerization. Using this approach, we found that this antibody was able to induce greater transcription of both *Il6* and *Lif* in Col14-LMC when added to Treg cell/Col14-LMC co-cultures, compared to IgG controls (**Fig. 5L**). Similar to IL-18, activation via 4-1BB (using this same agonistic antibody) has previously been shown to decrease immunosuppressive activity by Treg cells [47, 48]. Thus, 4-1BB stimulation seems to specifically induce repair activity in Treg cells, rather than non-specifically inducing all functional activities.

## Discussion

Transcriptomic and functional analysis of “tissue repair”-oriented Areg-producing immune cells has previously been unavailable due to the inability to sort live Areg-producing cells. Our novel *Areg^Thy1.1^* reporter mouse addresses this deficiency, which when paired with single cell RNA and TCR sequencing allowed us to interrogate reparative Treg cells in the lung in different models of damage at multiple timepoints of disease progression.

We found that our bulk RNA-seq efforts to elucidate differences in reparative Treg cells in models of lung damage were hampered by heterogeneity between sorted Thy1.1^−^ vs. Thy1.1^+^ cells, possibly related to the inclusion of different amounts of proliferating cells. Our use of scRNA-seq to address this heterogeneity revealed several different tissue subsets of Treg cells in the setting of lung damage. Certain subgroups, when compared with the Treg cells present in the lung in control mice, were only substantially detected during lung damage – the Ccr8 and Rorc subgroups identified here. Limiting our comparison to only these “induced” subsets of Treg cells allowed us to ascertain a defined gene signature associated with reparative, Areg-producing Treg cells. Several of the pathways identified in reparative Treg cells have not been previously studied in relation to tissue repair. For instance, Hedgehog signaling has been shown to mediate the conversion of Treg cells to a Th17-like phenotype in the context of breast cancer [49]. Whether such alterations are present and meaningful in the context of tissue damage remains to be seen; this study provides the groundwork for several such lines of investigation.

There is currently a surfeit of knowledge in the field of immunology regarding the specific antigens or other tissue cues that evoke expansion of T cells in the context of sterile damage at tissue sites. Our work here shows that Treg cells undergo progressive clonal expansion in the context of sterile lung injury (bleomycin). This may relate to increased access of reparative Treg cells to certain tissue self-antigens that are exposed upon lung damage. Alternatively, the lung microbiota, while known to be significantly less established than microbiota in other organs such as the gastrointestinal tract, is known to have an impact on the bleomycin model of ALI [50], and Treg cell expansion in this context could be in response to the microbiome. The fact that a group in a separate facility, which is likely to have a dissimilar lung microbiome, uncovered similar expanded clones to ones found here in a different model of lung fibrosis [36] would seem to indicate that a damage-induced self-antigen is a more likely candidate, but further studies (potentially with germ-free mice) must be done in this regard to fully address this point.

The potential antigen specificity of the clonally expanded Treg cells found in this study was not investigated here, but recent work has offered insight in this regard. One non-peer-reviewed study [41] (a doctoral thesis) found similar expanded TCR clones specifically on Treg cells in an immunosuppression-biased lung adenocarcinoma model. The two common clones that they select for further investigation, after finding them across lungs of multiple tumor-bearing mice, are two of the same ones we see in our shared/expanded set in bleomycin 21 dpi mice. This group performed yeast display peptide-MHC library experiments in an attempt to find antigens responsible for their expansion, but were unable to enrich for any specific target, concluding that these TCRs are potentially not specific to the purported MHCII subtype of this TCR, or have very low affinity to cognate antigen. In addition, they transduced T cells with one of these TCRs and adoptively transfer them to *Rag2*^−/-^ mice, but they failed to see greater localization to the lung, either at baseline or in a model of lung cancer. However, they suggest that other models of tissue damage may be better able to expose self-antigens to which these cells may be responding. A fully transgenic model encoding one of the most expanded TCRs from our dataset may be helpful in elucidating the effect of these common lung-expanded clones across models [7, 51].

Treg cell costimulation by the Tnf receptor superfamily (Tnfrsf genes) is known to have several important functions in activation and proliferation [52], but its effects on initiation of reparative activity in Treg cells is unknown. Our data show that in an in vitro assay for tissue repair activity, activation through 4-1BB heightens Treg cell signaling to lung mesenchymal target cells. Furthermore, this does not seem to be related to general activation of Treg cells when stimulated through this pathway, given that other work has shown that Treg cells are in fact less immunosuppressive when stimulated through this pathway [47, 48]. Given that other T cells can express 4-1BB [48], models wherein reparative effects of 4-1BB stimulation can be isolated specifically to Treg cells in vivo would be important for further establishing a role of signaling through this pathway in tissue repair.

In conclusion, the findings from this study provide a foundation for further investigation of Treg cell phenotypes and clonal expansion states associated with tissue reparative activity. Furthermore, the novel *Areg^Thy1.1^*knock-in mouse generated for this report could be used by other groups to investigate other aspects of reparative Treg cell biology, or other tissue cell types that produce Areg. There are many potential therapeutic benefits to harnessing optimal tissue repair abilities by Treg cells in the lung and at other tissue sites [53]; we hope that this work has offered some insight into the transcriptional and cellular pathways used by reparative Treg cells, that may ultimately assist in the development of these types of interventions.

## Methods

### Sex as a biological variable

For the transcriptomic studies and lung disease induction comparisons in this report, only male mice were used; this was done based on prior work indicating that Areg-mediated repair during influenza-induced lung damage in mice shows differences between sexes [54], and that bleomycin-induced lung damage is more pronounced in males [55]. All other experiments in this report utilize both male and female mice.

### Mice

*Areg^Thy1.1/Thy1.1^* mice were a novel creation for these studies. To create this strain, we inserted a P2A self-cleaving viral peptide followed by a mouse *Thy1^a^* (*Thy1.1*) sequence into the endogenous mouse *Areg* locus prior to the native stop codon with a subsequent FRT-flanked neomycin resistance cassette. Following recombineering of this construct and transduction, mouse embryonic stem cells positive for this construct were isolated by neomycin treatment, then microinjected into mouse embryos to create heterozygotes (on the C57BL6/N *Thy1^b^* [*Thy1.2*] background), and the FRT-flanked neomycin cassette was then removed by crossing with the FLPR mouse. We then homozygosed these mice by breeding to create *Areg^Thy1.1^*^/*Thy1.1*^ mice (referred to as *Areg^Thy1.1^* mice in the report). *Foxp3^GFP^*mice were a generous gift from the laboratory of Dr. Alexander Rudensky (Memorial Sloan Kettering), and were previously described [56]. Wild type (WT) mice (C57BL/6N) were acquired and bred from Jackson Laboratory stocks (Strain #:005304). These mice or lab-bred descendants were utilized for Col14-LMC isolation/sorting.

### Bulk RNA-seq

Treg cells were harvested and prepared for flow cytometry as described, then sorted from 4 IAV-treated mice (275 TCID50 PR8/H1N1) (8 dpi) and 3 bleomycin-treated mice (1 U/kg) (12 dpi); all mice were *Foxp3^GFP^Areg^Thy1.1^*. 5,000 Thy1.1^−^ and 5,000 Thy1.1^+^ cells were sorted per sample. Following sorting, samples were centrifuged at 550 x g/8 min./4°C, supernatants were aspirated, and samples flash frozen. RNA was extracted, cDNA libraries were generated, and sequencing was performed by Genewiz from Azenta Life Sciences. Fastq files were aligned using RNA Detector [57], with adapter trimming using Trim Galore and pseudoalignment using Salmon to generate RNA counts. DESeq2 (in the R interface) was used for analysis (paired analysis between Thy1.1^−^ and Thy1.1^+^ cells from each mouse). ComplexHeatmap [58] and EnhancedVolcano R packages were used for analysis, and GSEA [59] was used for pathway analysis. Venny was used for Venn diagram analysis.

### Single cell RNA– and TCR-seq

For the first experiment, 7 IAV-treated mice (275 TCID50 PR8/H1N1) (5 dpi), 5 bleomycin-treated mice (1 U/kg) (12 dpi), and 2 control mice (saline-treated, 5 dpi) were used; for the second experiment, 5 bleomycin-treated mice (1 U/kg) (21 dpi), and 2 control mice (saline-treated, 5 dpi) were used. All mice were *Foxp3^GFP^Areg^Thy1.1^*. Treg cells were harvested from mouse lungs and prepared for flow cytometry staining as described. Cells from separate mice were stained for hashes, stained for sorting antibodies, and sorted. Hashing was performed with custom αCD2 hashes (10 min. of FC block incubation prior to staining, 0.5 ug of hash per 2 million cells for 30 min. at 4°C); custom hashes were conjugated as described in protocols of the Technology Innovation Lab at the New York Genome Center using iEDDA-click chemistry. Afterwards, cells from separate mice were combined, stained for sorting antibodies, and sorted on a BD Aria sorted. Cells amounts to sort were determined based on optimal protocols for 10x Chromium Beads (∼1000-3000 cells per mouse; ∼15,000 total cells per bead). Thy1.1^−^ and Thy1.1^+^ cells were run on separate beads in parallel to eliminate the need for a separate hashing step post-sorting. Thy1.1^−^ and Thy1.1^+^ Treg cells were included in roughly equal amounts, to give us the ability to better determine heterogeneity and representation within the Thy1.1^+^ group. Control mouse Treg cells were all Thy1.1^−^. Sorted Treg cells were loaded as described by the manufacturer protocols onto 10x Chromium Genomic V1 Platform (10x Genomics) and libraries were prepared for single cell 5’ transcripts, TCRαβ transcripts, and CITE-seq hash libraries (Chromium Single Cell V(D)J Reagent Kit). SPRIselect Beads (BD) were used in the preparation process. Cell Ranger software (v5.0.0) (10x Genomics) was used to process reads (gene expression, CITE-seq, TCRαβ), with raw base call files demultiplexed with cellranger mkfastq function. This was subsequently aligned to the mm10 reference genome, then unique molecular identifiers (UMI) were collapsed to count matrices using cellranger multi function. Raw hash reads were processed for analysis with the CITE-seq-Count (GitHub), in order to create a hash count matrix. Hashes were then normalized by total counts across cells, with total UMI counts set to 100, with identity assigned to the most highly expressed hashtag that represented a minimum of 2/3 of the recovered mapped UMIs in a cell. Cells with suboptimal UMI total counts (UMI total < 5 or UMI total > 5) were removed from downstream analyses. The Seurat package (in R) [19] was used for analysis of single cell datasets. Cells were subsetted to exclude cells with < 200 or > 3500 features or a mitochondrial percent of features > 5%. Top 2000 top variable features were used to determine clustering (with TCR genes excluded). Since Thy1.1^−^ and Thy1.1^+^ cells were run on separate beads, Combined Cluster Analysis (CCA) was performed to adjust for any batch effects between samples. 20-25 principal components were used to generate UMAPs, with clustering resolution set from 0.7-1.9 for different datasets. “Pseudobulking” of expression values for each mouse was used for certain analyses. EnhancedVolcano R package, Immunarch R pacakage, ggPlot2 R package, and Upset plots [60] were used for analyses.

### Additional methods

For detailed methods regarding “Mouse lung damage models and assessment”, “Lung, spleen, and lymph node processing”, “Splenic/lymph node Treg cell stimulation protocols”, “CD4 T cell bead enrichment and Treg cell sorting”, “Flow cytometry”, “Col14-LMC negative enrichment and sorting”, “Col14-LMC/Treg cell co-culture”, and “RNA extraction and qPCR”, see Supplementary Methods.

### Statistics

R and GraphPad Prism (v10.1.2) was used for all statistical analyses and graphing. For flow cytometry, qPCR, or TCR diversity (Chao1) analysis where two groups were compared, two-tailed unpaired Student’s t-tests were used. For longitudinal weight loss, body temperature, or SpO_2_ analysis, two-way repeated measure ANOVA was used. Statistical significance was determined at p<0.05, with further levels of significance reported in figure legends. Sample size estimation was determined based on previous studies. FlowJo (v10), Microsoft Excel, Microsoft Powerpoint, SnapGene Viewer, Adobe Illustrator, MacVector, and R were used to set up experiments, analyze data, and prepare data.

### Study approval

Animal experiments were approved by Columbia University’s Institutional Animal Care and Use Committee (protocol AC-AABT2656).

### Data availability

Sequencing data associated with this manuscript have been deposited in NCBI’s Gene Expression Omnibus (GEO Accession Numbers: GSE277226 and GSE277256). We report no original source code from this manuscript. All other data have been included in the manuscript or supplemental information.

## Acknowledgements

We thank several Columbia University Core Facilities for their contributions to this work: the Herbert Irving Cancer Comprehensive Cancer Center Genetically Modified Mouse Models Core Facility, the Microbiology & Immunology Shared Resources, Columbia Stem Cell Initiative (CSCI) Flow Cytometry, and the Herbert Irving Cancer Comprehensive Cancer Center Molecular Pathology Shared Resource (MPSR), funded in part through the NIH/NCI Cancer Center Support Grant P30CA013696. We thank Dr. Alexander Rudensky for contribution of *Foxp3^GFP^* mice. We thank Dr. Donna Farber for contribution of influenza A virus (PR8/H1N1). We thank all other Arpaia lab members for their feedback and comments on this work. This work was supported by NIH grant R01HL148718.

## Author contributions

L.F.L., K.A.K., A.H., and N.A. designed research; L.F.L., K.A.K., A.K., S.R., and K.d.l.S.-A. performed research; L.F.L., K.A.K., A.K., and S.R. analyzed data; L.F.L., K.A.K., and N.A. wrote the paper. Order of co-first authorship was determined as such due to final assembly of manuscript by L.F.L.

## Competing interests

The authors claim no competing interests.

---

**Figure S1.**
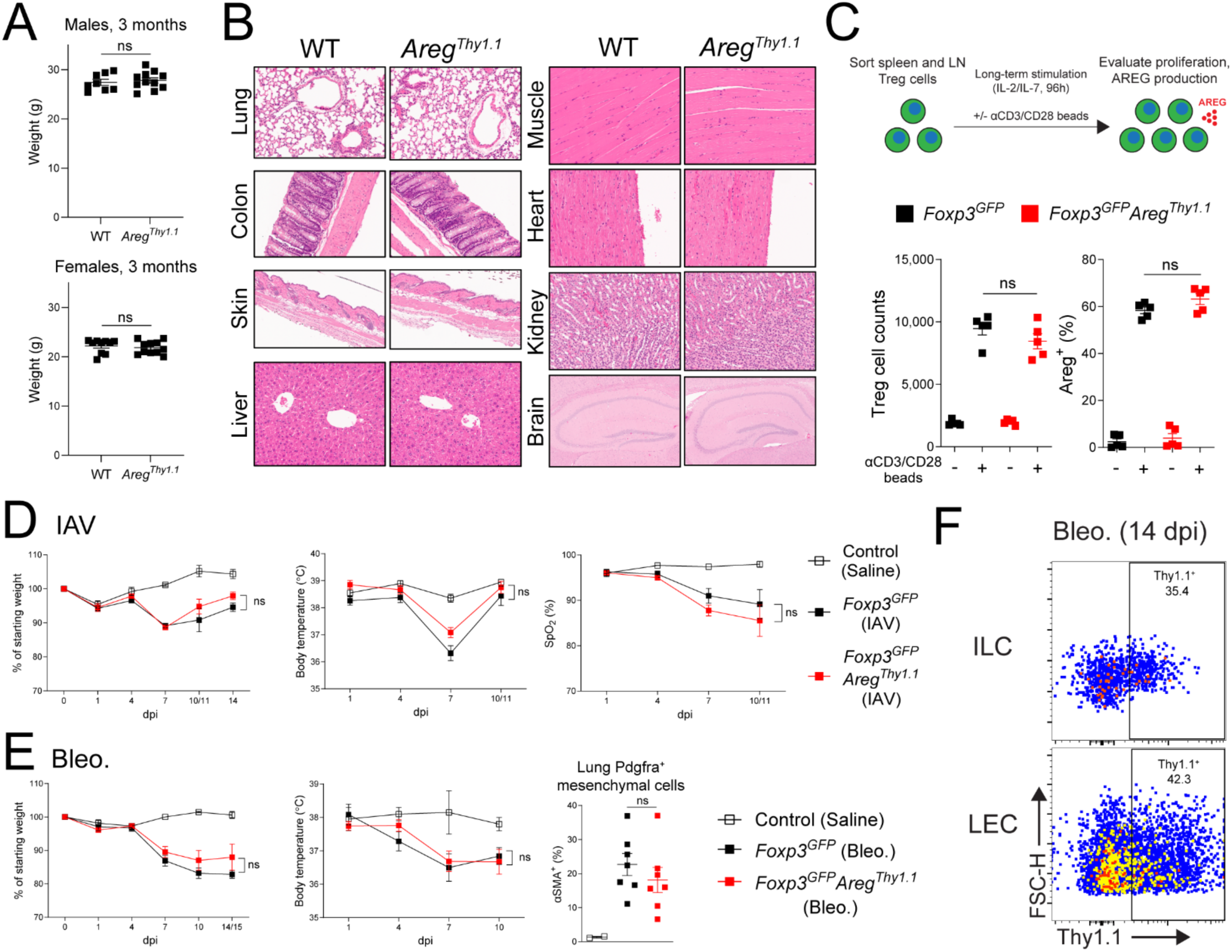
Supplementary data from Figure 1. **A)** Weights of WT or *Areg^Thy1.1^* mice (male or female) at 3 months of age. From 2 separate cohorts of mice of each genotype. **B)** Histology from indicated organs of WT or *Areg^Thy1.1^* mice (hematoxylin & eosin staining). All sections are at 20x magnification besides the brain (5x). Representative histology shown from 2 separate experiments. **C)** Treg cells were sorted from spleen/lymph node cell suspensions from *Foxp3^GFP^*or *Foxp3^GFP^Areg^Thy1.1^* mice, then cultured for 96h with IL-2 and IL-7, with or without the inclusion of αCD3/CD28 beads (see schematic). Following this, Treg cells were counted and stained for endogenous AREG protein production. Graph contains all values from 2 separate experiments. **D)** *Foxp3^GFP^* or *Foxp3^GFP^Areg^Thy1.1^* mice were treated with IAV (PR8-H1N1, 100 TCID50, intranasal), with weight, body temperature, and blood oxygen saturation measured every 3-4 days to assess course of disease. n=5-6 mice per IAV groups, 2 mice in control group. Graphs contain all values from 2 separate experiments. dpi: days post-instillation. **E)** *Foxp3^GFP^* or *Foxp3^GFP^Areg^Thy1.1^* mice were treated with bleomycin (1 mg/kg, intratracheal), with weight, and body temperature measured every 3-4 days to assess course of disease. To assess fibrosis as a functional readout for this model, lungs at terminal timepoint (14-15 dpi) were processed for flow cytometry, wherein Pdgfra^+^ mesenchymal cells were intracellularly stained for expression of smooth muscle actin (αSMA), a proxy for fibrosis induction. n=7 mice per bleomycin groups, 2 mice in control group. Graphs contain all values from 2 separate experiments. **F)** Staining for Thy1.1 in ILC and LEC in the lungs of *Foxp3^GFP^Areg^Thy1.1^*mice undergoing the bleomycin model (14 dpi). Representative staining from 3-4 experiments. See Fig. S2A for gating strategies throughout figure. Standard error displayed on graphs; n.s: not significant, *: 0.01<p<0.05, **: 0.001<p<0.01, ***: 0.0001<p<0.001, ****: p<0.0001.

**Figure S2.**
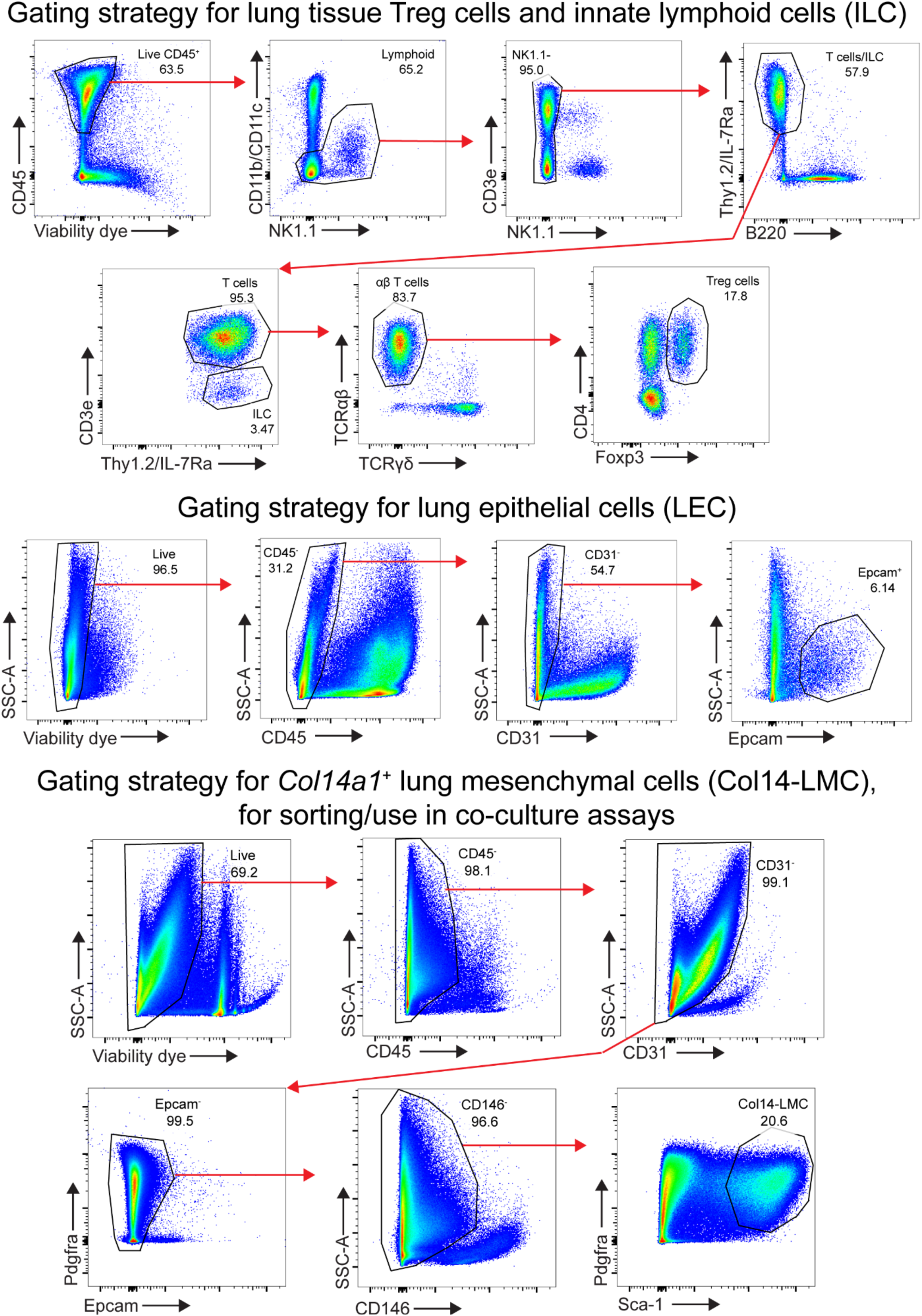
Gating strategies used in experiments. Flow cytometric gating strategies for assessing lung tissue Treg cell, lung tissue innate lymphoid cell (ILC), and lung epithelial cells (LEC), as well as that used for sorting Col14-LMC reparative mesenchymal cells.

**Figure S3.**
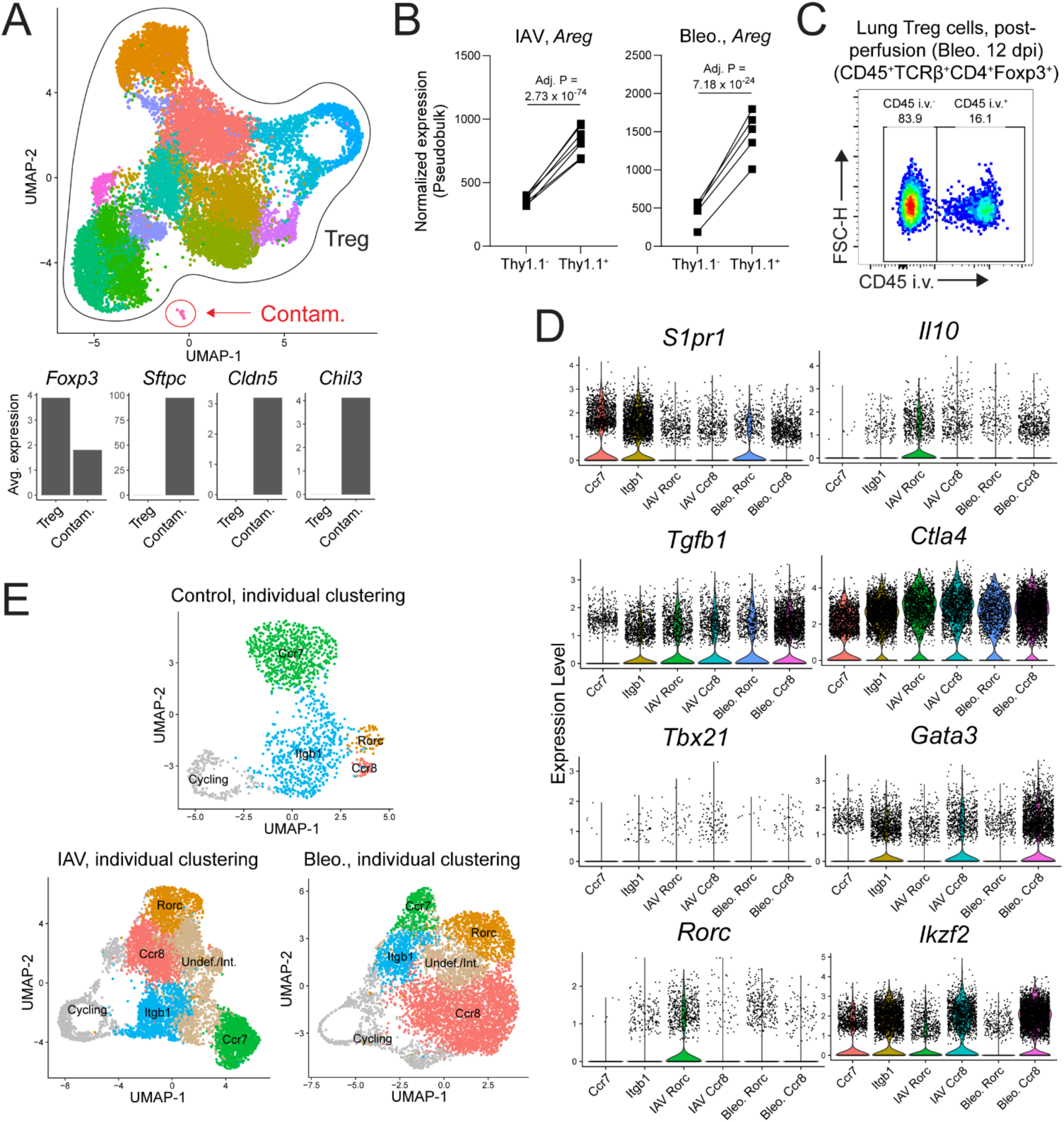
Supplementary data from Figure 3. **A)** Top: UMAP of clustered cells from sc gene expression analysis using Seurat (see Fig. 3A), prior to removal of contaminating non-Treg cells (“Contam.” on graph). Bottom: Graphs indicating average expression of *Foxp3* (Treg cell-specific) or Treg cell-nonspecific genes (*Sftpc*: epithelial, *Cldn5*: endothelial, *Chil3*: macrophage), in contaminating population compared to other cells in dataset (“Treg”). **B)** *Areg* expression from separate mice (counts aggregated and normalized via pseudobulk analysis), with paired analysis of Thy1.1^−^ vs. Thy1.1^+^ Treg cells, from IAV and bleomycin datasets. **C)** Flow cytometry of Treg cells from bleomycin-treated lungs (14 dpi), treated intravenously prior to harvest with CD45 antibody (“CD45 i.v.”) to identify circulating cells. Lungs had undergone an identical perfusion procedure to that used for all experiments herein. Representative staining from 3 experiments. **D)** Violin plots indicating per-cell expression of select genes in combined control/IAV/bleomycin dataset. **E)** Individual re-clustering of control, IAV, and bleomycin cells for better delineation of subgroups.

**Figure S4.**
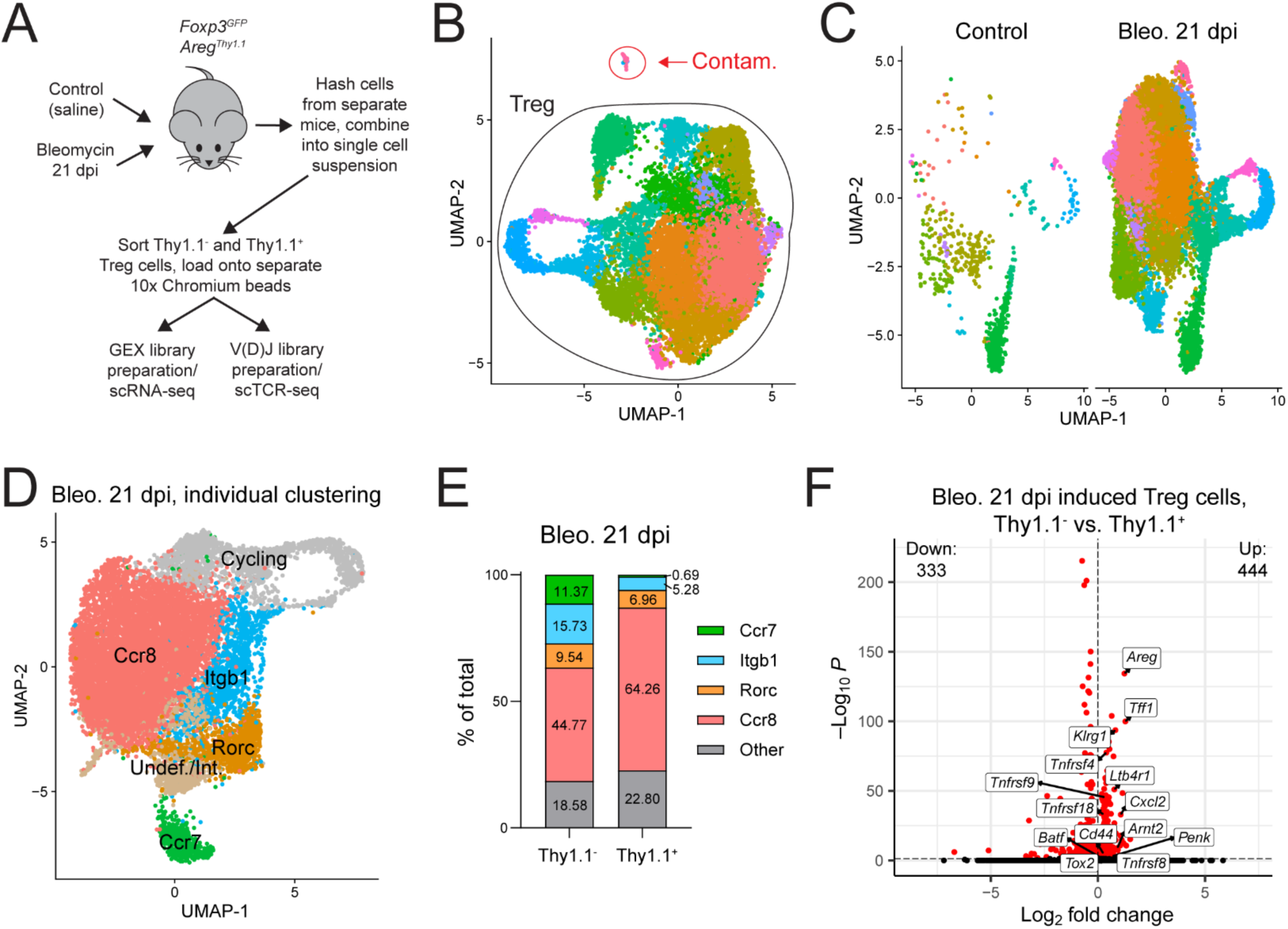
scRNA-seq of Areg-producing and non-producing lung Treg cells from bleomycin-treated mice at a later timepoint. **A)** Schematic of single cell (sc) RNA-seq experiments using lung Thy1.1^−^ vs. Thy1.1^+^ Treg cells from the bleomycin model at 21 dpi (as well as control, saline-treated mice). **B)** UMAP of clustered cells from sc gene expression analysis of the bleomycin 21 dpi dataset using Seurat, prior to removal of contaminating non-Treg cells (“Contam.” on graph). **C)** UMAP of cell clustering after removal of contaminating cells, split by treatment status of Treg cells. **D)** Individual re-clustering of bleomycin 21 dpi cells for better delineation of subgroups. **E)** Proportions of each assigned subgroup in Thy1.1^−^ and Thy1.1^+^ Treg cells from the bleomycin 21 dpi dataset (from individual clustering). **F)** Volcano plot of DEGs from Thy1.1^−^ vs. Thy1.1^+^ induced Treg cells (Ccr8 and Rorc subgroups combined) from bleomycin 21 dpi dataset. Red dots on volcano plots: significant DEGs (FDR adj. p-value < 0.05). No fold change cutoff. Numbers of significantly upregulated and downregulated genes indicated on plots.

## Supplementary Methods

### Mouse lung damage models and assessment

Influenza A (IAV) virus (PR8/H1N1) was a generous gift from the laboratory of Dr. Donna Farber (Columbia University). For IAV infection, mice were given ketamine/xylazine for anesthesia, then infected intranasally with 100-300 TCID50 of virus diluted in 1x PBS (as determined by the Farber lab using Madin-Darby canine kidney epithelial cell infection assays). 10-16 week old male mice were used for IAV experiments, and mouse lungs were harvested at 5 or 8 days post-inoculation (dpi). Bleomycin (Sigma or Teva) was diluted in sterile 0.9% saline (0.5 U/ml). Bleomycin was administered to mice using one of two methods for different experiments: surgical intratracheal or oropharyngeal intratracheal administration. For surgical intratracheal administration, we followed a previously described technique (Orlando et al. 2019). Briefly, mice were given buprenorphine prior to experimentation for analgesia, then ketamine/xylazine for anesthesia. Once unconscious and unresponsive to toe pinch, the area of surgery was cleaned using 3 alternating scrubs of PVP Iodine Prep Pads (Medline) and WEBCOL Alcohol Preps (Covidien). A scalpel was then used to make an incision in the center of the neck, then the salivary glands were split and the muscles overlaying the trachea were carefully cut to expose the trachea. An insulin syringe (Corning) was loaded with 30-50 µl of bleomycin (calculated for 1 U/kg administration), with air pockets above and below liquid to distribute liquid into lungs. The syringe was inserted into the trachea and plunged to aspirate the solution. The surgical incision was sealed with Vetbond Tissue Adhesive (3M) and reinforced with Reflex 9mm Wound Clips (Roboz Surgical Instrument). Buprenorphine was administered 3 times at 12h intervals following surgery for analgesia, and mice were monitored for 10 days to ascertain wound closure and healing. For oropharyngeal intratracheal administration, 50 µl per mouse was administered oropharyngeally (∼1 U/kg), following a previously described technique (De Vooght et al. 2009). Briefly, mice were given ketamine/xylazine for anesthesia, then placed on an apparatus suspending them at a 60° angle from horizontal by surgical suture string from their teeth. The tongue was removed from the mouth and held with padded forceps, then the bleomycin solution was pipetted into the back of the mouth, followed immediately by plugging the nose with padded forceps to induce oral inhalation of the solution. 10-16 week old male mice were used for bleomycin experiments, and mouse lungs were harvested at 11-21 dpi. Littermate, age-matched mice were used for all experiments. For IAV and bleomycin experiments, mice were weighed every 1-4 days. In certain experiments, mouse body temperature was assessed with a rectal thermometer every 1-4 days. For IAV experiments, mouse blood oxygen saturation (SpO_2_) was assessed using a MouseOx Plus Pulse Oximeter (Starr Life Sciences). The area around mouse the mouse neck was shaved and Nair Hair Remover Product was applied to chemically remove residual hair at the time of IAV infection. A Small Mouse Collar Sensor (Starr Life Sciences) was used to assess SpO_2_ on unanesthetized at indicated timepoints during IAV infection progression, with mice placed in a 1 L beaker to restrict movement; assessment was taken for 2-5 min. per mouse at each timepoint, with only high-quality, error-free readings taken and averaged to determine a composite SpO_2_ value. For histology of various organs, lung, colon, skin, liver, muscle, heart, kidney, and brain were isolated from untreated mice without perfusion, fixed in 10% neutral buffered formalin (Epredia), then sent to Histology Consultation Services for embedding in paraffin, sectioning at 5 µm, and H&E staining; full organs were imaged using a Aperio AT2 (Leica) full slide scanner.

### Lung, spleen, and lymph node processing

For lung processing, mice were euthanized and dissected to expose the lungs. Perfusion of the lungs was performed, after nicking the left femoral artery and the left atrium of the heart, through the left ventricle of the heart with 10 ml of cold 1x PBS. Lungs were excised and placed in 0.5 ml tissue preparation media (RPMI with 100x penicillin/streptomycin, 100x GlutaMAX, 100x HEPES [all Gibco], and 5% FBS [Corning]) in a 5 ml Eppendorf tube, where they were minced. 3.5 ml was added to tubes of tissue preparation media with 5 U/ml DNAse, 1 mg/ml of collagenase A, and 1 mg/ml of dispase. (Dispase was pre-dissolved in 1x PBS consisting of 10% of the volume of the final digestion mixture.) Lungs were digested in shaking incubator set to 110 rpm at 37°C for 1h. Suspensions were then poured over a 100 µm cell strainer (Corning) into a 50 ml Falcon tube (Corning), pushed through the mesh with the top of a syringe, rinsed with 10 ml tissue preparation media, pushed through again, then rinsed with 5 ml tissue preparation media. Cells were centrifuged at 450 x g/4°C/5 min., then supernatants were poured off (due to delicate nature of pellet resulting from use of dispase). Pellets were resuspended in 2 ml of 1x ACK lysis buffer (deionized water with 154 mM ammonium chloride [Fisher], 10 mM potassium bicarbonate [Fisher], and 0.1 mM ethylenediaminetetraacetic acid [EDTA] disodium salt dihydrate [Fisher], pH 7.2) and incubated at room temperature for 2 min., then quenched with 10 ml tissue preparation media and ran through a 100 µm nylon mesh sheet into a 15 ml Falcon tube (Corning). Cells were centrifuged at 450 x g/4°C/5 min., the supernatant was aspirated, and cells were resuspended in 1 ml of tissue preparation media and placed on ice. Cells were then used for bead enrichment/sorting or antibody staining for flow cytometry. For spleen and lymph node processing, mice were euthanized and dissected to extract the spleen (and in some experiments, inguinal and cervical lymph nodes as well), which were excised and placed on top of a 100 µm cell strainer (Corning) in 5 ml of tissue preparation media in a 6 cm dish (Corning), pushed through the mesh with the top of a syringe, rinsed with 5 ml tissue preparation media, pushed through again, then rinsed with 5 ml tissue preparation media; contents of the dish were placed in a 15 ml Falcon tube (Corning). Cells were centrifuged at 450 x g/4°C/5 min., then supernatants were aspirated. Cells were then resuspended in 2 ml of 1x ACK lysis buffer and incubated at room temperature for 2 min., then quenched with 10 ml tissue preparation media and ran through a 100 µm nylon mesh sheet into a 15 ml Falcon tube (Corning). Cells were centrifuged at 450 x g/4°C/5 min., the supernatant was aspirated, and cells were resuspended in 1 ml of tissue preparation media and placed on ice. Cells were then used for bead enrichment/sorting, or directly in short-term stimulation experiments.

### Splenic/lymph node Treg cell stimulation protocols

For the short-term stimulation protocol, mouse splenocytes were isolated as described in “Lung, spleen, and lymph node processing”, then cultured (bulk splenocytes, unsorted) for 3h in complete T cell media (RPMI + 100x penicillin/streptomycin, 100x GlutaMAX, 100x HEPES, 100x sodium pyruvate, 100x nonessential amino acids, 1000x β-mercaptoethanol [all Gibco], and 10% fetal bovine serum [FBS]) with phorbol 12-myristate 13-acetate (PMA) (50 ng/ml) (Sigma) and ionomycin (500 ng/ml) (Sigma); cells were then stained for flow cytometry to identify Treg cells. For the long-term stimulation protocol, mouse spleen and lymph node T cells were isolated as described in “Lung, spleen, and lymph node processing”, then negatively enriched for CD4 T cells and sorted for Treg cells as described in “CD4 T cell bead enrichment and Treg cell sorting”. Treg cells were then cultured in complete T cell media for 96h in round bottom plates in the presence of rhIL-2 (200 U/ml; NCI Preclinical Repository) and rhIL-7 (10 ng/ml; NCI Preclinical Repository), with or without Dynabeads Mouse T-Activator CD3/CD28 for T-Cell Expansion and Activation (Thermo) beads added at a 1:1 Treg cell/bead ratio. Cells were then counted using 123count eBeads Couting Beads (Invitrogen) on a BD LSRII. Cells were also stained for AREG production using the endogenous AREG antibody (see “Flow cytometry”).

### CD4 T cell bead enrichment and Treg cell sorting

Lungs, spleens, and/or lymph nodes were prepared as described in “Lung, spleen, and lymph node processing”. Lung, spleen, and/or lymph node cells were then enriched for CD4 T cells using either positive enrichment with the Dynabeads FlowComp Mouse CD4 kit, or negative enrichment with iMag Streptavidin Particles Plus – DM beads (BD), according to their respective protocols. For negative enrichment, following a 10 min. incubation with FC block (purified anti-mouse CD16/CD32, clone 2.4G2; Cytek), cells were incubated in a cocktail of biotinylated antibodies towards mouse TER-119 (clone TER-119; Biolegend), CD31 (clone 390; Biolegend), Epcam (clone G8.8; Biolegend), Pdgfra (clone APA5; Biolegend), CD19 (clone 6D5; Biolegend), NK1.1 (clone PK136; Biolegend), CD11b (clone M1/70; Biolegend), CD11c (clone N418; Biolegend), and CD8a (clone 53-6.7; Biolegend) (only TER-119, CD19, NK1.1, CD11b, CD11c, and CD8a antibodies were used for spleen/lymph node preparations). Post-bead enrichment, cells were stained with fluorescent mouse antibodies for flow cytometric sorting (100 µl per mouse). Antibodies used were: CD45-BV786 (clone 30-F11; BD), CD11b-BV510 (clone M1/70; BD), CD11c-BV510 (clone HL3; BD), TCR β-BV711 (clone H57-597; BD), CD45R (B220)-PerCP-Cy5.5 (clone RA3-6B2; Cytek), CD8a-PE (clone 53-6.7; Cytek), NK1.1-PE-Cy7 (clone PK136; Biolegend), CD4-BV605 (clone RM4-5; BD), and CD90.1 (Thy1.1)-APC (HIS51; Invitrogen). Treg cell preparations were done with lungs of *Foxp3*^EGFP^ mice, (i.e., Treg cells express GFP), allowing sorting for GFP^+^ cells. Cells were incubated with antibodies for 20 min. at 4°C. Post-staining, cells were washed, then resuspended in 200 µl per mouse of FACS buffer without sodium azide, ran through a 40 µm mesh, then sorted on a BD Aria sorter, with Sytox Blue (Invitrogen) added 5 min. prior to running samples for dead cell exclusion.

### Flow cytometry

Lungs were prepared for flow cytometry as described above, with 2-3 million cells from final single cell suspensions used for antibody staining and 100,000-1 million cells ran on a BD LSRII or BD Fortessa. Cells were stained in flow cytometry buffer (1x PBS with 1% BSA [Gold Biotechnology], 2.5 mM ethylenediaminetetraacetic acid [EDTA] disodium salt dihydrate [Fisher], and 0.1% sodium azide [Fisher]). Zombie Violet Fixable Viability Kit (Biolegend) or GhostDye Red 780 (Cytek), stained in a separate step from surface antibodies in 1x PBS, or Sytox Blue (Thermo), added directly to sample 5 min. prior to running on cytometer, were used for dead cell exclusion. Staining was preceded by a 10 min. incubation with FC block (purified anti-mouse CD16/CD32, clone 2.4G2; Cytek) in flow cytometry buffer, with 2x antibody cocktail added. For analysis of Treg cell/innate lymphoid cell (ILC) populations from lung single cell suspensions, surface staining was done using these anti-mouse antibodies: NK1.1-BUV395 (clone PK136; BD), CD45-BUV395 (clone 30-F11; BD), CD3e-BUV496 (clone 145-2C11; BD), CD4-BUV737 (clone RM4-5; BD), TCR γ/δ-BV421 (clone GL3; Biolegend), CD11b-BV510 (clone M1/70; BD), CD11c-BV510 (clone HL3; BD), CD4-BV510 (clone RM4-5; Biolegend), TCR β-BV711 (clone H57-597; BD), CD45-BV786 (clone 30-F11; BD), CD45R (B220)-PE (clone RA3-6B2; Cytek), TCR β-PE/Dazzle594 (clone H57-597; Biolegend), CD90.2 (Thy1.2)-PE-Cy7 (53-2.1; Biolegend), CD127 (IL-7Ra)-PE-Cy7 (A7R34; Cytek), NK1.1-PE-Cy7 (clone PK136; Biolegend), CD3e-PE-Cy7 (clone 145-2C11; Cytek), and/or CD4-APC-Cy7 (clone RM4-5; Cytek). For analysis of lung epithelial cells (LEC) from lung single cell suspensions, surface staining was done using these anti-mouse antibodies: CD45-BUV395 (clone 30-F11; BD), CD31-BV605 (clone 390; Biolegend), and Epcam-PerCP-Cy5.5 (clone G8.8; Biolegend). For analysis of smooth muscle actin on Pdgfra^+^ mesenchymal cells from lung single cell suspensions, surface staining was done using these anti-mouse antibodies: CD45-BUV395 (clone 30-F11; BD), CD31-BV711 (clone 390; Biolegend), Epcam-BV785 (clone G8.8; Biolegend); CD146-PerCPCy6.6 (clone ME-9F1; Biolegend), and Pdgfra-BV605 (clone APA5; Biolegend). For analysis of Thy1.1 on cell populations (live cells), anti-mouse CD90.1 (Thy1.1)-APC (HIS51; Invitrogen) antibody was used. Following surface marker staining, for staining with anti-mouse Foxp3-FITC (clone FJK-16s; Invitrogen), anti-mouse Ki67-AlexaFluor700 (clone SolA15; Invitrogen), anti-mouse α-smooth muscle actin-eFluor660 (clone 1A4; Invitrogen), or biotinylated anti-mouse AREG (endogenous) (polyclonal goat IgG; R&D systems) followed by Streptavidin conjugates, cells were fixed/permeabilized with Foxp3/Transcription Factor Staining Buffer Kit (Cytek). For analysis of endogenous AREG on cell populations, biotinylated anti-mouse AREG (polyclonal goat IgG; R&D Systems) was used, followed by a separate staining step with Streptavidin-PE-eFluor610 (Invitrogen) or Streptavidin-PE (Thermo). For Treg cells from bleomycin-treated mouse lungs (live cells used for staining), proteins were assessed with flow cytometry using these antibodies: anti-mouse OX-40-PE (clone OX-86; Biolegend), anti-mouse 4-1BB-PE (clone 17B5; Biolegend), anti-mouse CD51-PE (clone RMV-7; Biolegend), and anti-human/mouse Ltb4r1 (clone 7A8; Sigma); since anti-mouse/human Ltb4r1 is an unconjugated mouse monoclonal antibody, the Mouse-on-Mouse Immunodetection Kit (Vector Laboratories) was used (including biotinylated anti-mouse IgG), with subsequent staining using Streptavidin-PE (Thermo). Fluorescence-minus-one (FMO) controls were included to define staining boundaries where necessary. For certain experiments, CD45-APC (clone 30-F11; Cytek) was injected intravenously into mice 5 min. prior to euthanasia, to label circulating immune cells.

### Col14-LMC negative enrichment and sorting

Lungs were prepared as described in “Lung processing”. Lung cells were then negatively enriched for mesenchymal cells using iMag Streptavidin Particles Plus – DM beads (BD), according to their protocol. For negative enrichment, following a 10 min. incubation with FC block (purified anti-mouse CD16/CD32, clone 2.4G2; Cytek), cells were incubated in a cocktail of biotinylated antibodies towards mouse CD45 (clone 30-F11; Biolegend), CD31 (clone 390; Biolegend), Epcam (clone G8.8; Biolegend), and TER-119 (clone TER-119; Biolegend). Post-bead enrichment, cells were stained with fluorescent mouse antibodies for flow cytometric sorting (100 µl per mouse); antibodies used were: CD31-BV605 (clone 390; Biolegend), Epcam-PerCP-Cy5.5 (clone G8.8; Biolegend); Pdgfra-PE (clone APA5; Biolegend), CD146-PE-Cy7 (clone ME-9F1; Biolegend), CD45-APC (clone 30-F11; Cytek), and Sca-1-APC-Cy7 (clone D7; Biolegend). Post-staining, cells were washed, then resuspended in 200 µl per mouse, ran through a 70 µm mesh, then sorted on a BD Aria sorter, with Sytox Blue added 5 min. prior to running samples for dead cell exclusion.

### Col14-LMC/Treg cell co-culture

Col14-LMC were isolated/sorted as described and plated in mesenchymal cell media at 40,000-50,000 cells/well in 48 well tissue culture-treated plates (Corning). Cells were allowed to adhere overnight (16-18h); the following day, media was aspirated from cells, cells were washed with RPMI (Gibco) to remove dead cells, and fresh T cell media (RPMI + 100x penicillin/streptomycin, 100x GlutaMAX, 100x HEPES, 100x sodium pyruvate, 100x nonessential amino acids, 1000x β-mercaptoethanol [all Gibco], and 10% fetal bovine serum [FBS]) was added; cells were rested for ∼12h while Treg cell isolation/sorting occurred (∼8h). Treg cells from lungs or spleens of bleomycin-treated mice (11-15 dpi) were isolated/sorted as described above. 20,000-25,000 Treg cells (1:2 Treg cell:Col14-LMC ratio) were then added directly to Col14-LMC wells in small volumes (1/25 volume of media in wells). Dynabeads Mouse T-Activator CD3/CD28 for T-Cell Expansion and Activation (Thermo) beads were added to Treg cells at a 1:1 Treg cell/bead ratio prior to addition to wells. rhIL-2 (200 U/ml; NCI Preclinical Repository) and rhIL-7 (10 ng/ml; NCI Preclinical Repository) were added directly to wells at time of Treg cell addition. For some experiments, anti-mouse AREG antibody (polyclonal goat IgG; R&D Systems) (1 µg/ml) or normal goat IgG (R&D Systems) (1 µg/ml) were added to co-cultures. For some experiments, Treg cells were separated from Col14-LMC with a 0.4 µm pore size Transwell 24-well Permeable Membrane Insert (Corning). For some experiments, recombinant mouse IL-18 (Biolegend) (100 ng/ml) or vehicle was added to co-cultures. For some experiments, recombinant mouse 4-1BB ligand (Biolegend) (100 ng/ml), mouse vitronectin (Sino Biological) (100 ng/ml), and leukotriene B4 (Cayman) (100 ng/ml), or vehicle, were added to co-cultures. For some experiments, Ultra-Leaf Purified anti-mouse OX-40 agonistic antibody (clone OX-86; Biolegend) (10 µg/ml) or control Rat IgG1, κ (Biolegend) (10 µg/ml) was added to co-cultures. For some experiments, InVivoMAB anti-mouse 4-1BB agonistic antibody (clone 3H3; BioXCell) (10 µg/ml) or Rat IgG2a, κ Isotype Control (Biolegend) (10 µg/ml) was added to co-cultures. Co-cultures were incubated for 12h. At this time, wells were subjected to one wash of 5 mM EDTA in 1x PBS, and two additional washes of 1x PBS to remove Treg cells; following this removal, Col14-LMC in wells were lysed and analyzed for RNA (see “RNA extraction and qPCR”).

### RNA extraction and qPCR

RNA extraction was done using Trizol Reagent (Thermo) for lysis, followed by chloroform-based separation and precipitation of RNA with isopropanol. cDNA was created using a qScript cDNA Synthesis Kit (Quanta). qPCR was performed on cDNA using SYBR Green qPCR Master Mix (Thermo) and a Bio-Rad CFX384 qPCR system. Primers were created for this study using Integrated DNA Technology’s PrimerQuest platform, and University of California Santa Clara’s In-Silico PCR platform; all primers were ordered from Integrated DNA Technology and are listed in Supplementary Methods Table 1. Analysis was done by calculating ΔΔCt values relative to housekeeping gene (*Hprt*).

**Supplementary Methods Table 1.**
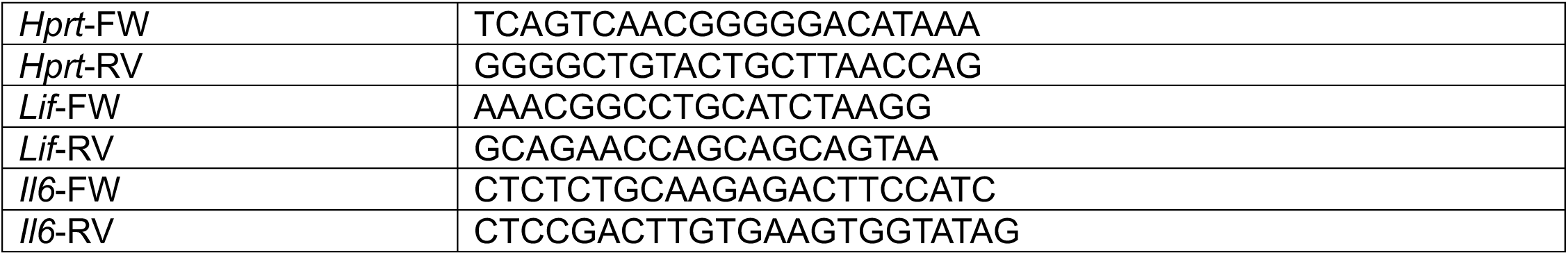
qPCR primers used in this study.

## Notes

### Competing Interest Statement

The authors have declared no competing interest.

